# LaCONIC: A Label-Aware and Graph-Guided Contrastive Multi-Omics Collaborative Learning Model for Cancer Risk Prediction

**DOI:** 10.1101/2025.11.26.690662

**Authors:** Pei Liu, Xiao Liang, Jiawei Luo

## Abstract

Investigating accurate cancer survival prediction models has important clinical value for optimizing therapeutic strategies and improving clinical outcomes. Although an increasing number of models have shifted from relying solely on clinical variables to integrating multi-omics data, there remains insufficient exploitation and integrative utilization of gene regulatory network (GRN) structures and cancer subtype information, which limits the predictive accuracy and biological interpretability of multi-omics models. Therefore, this study proposes label-aware and graph-guided multi-omics collaborative learning framework, termed LaCONIC, to achieve accurate and interpretable cancer survival prediction. To obtain biologically meaningful regulatory features of different molecular types, we first construct a gene regulatory network comprising multiple molecular entities and their relations, and design a heterogeneous graph self-supervised pre-training module to obtain unified graph-based gene representations. To fully leverage multiomics and multi-modal information, we develop an adaptive cross-omics representation learning module that performs intra-omics modality alignment (i.e., graph and expression modals) and inter-omics representation alignment, thereby achieving synergistic integration of multi-source information. Moreover, to comprehensively capture global cross-omics molecular interactions, we present a graph-guided cross-omics interactive learning module that explicitly encodes gene regulatory network priors within a Transformer-based architecture. Finally, we introduce cancer subtype information to construct a label-aware constraint mechanism that improves predictive performance while enhancing inter-class separability and intra-class consistency of the learned representations. Experiments on multiple cancer datasets show that LaCONIC outperforms existing state-of-the-art methods in various metrics. SHAP-based interpretability analysis on breast cancer further identifies survival-associated regulatory modules and high-risk molecules (e.g., hsa-miR-148a-3p, ARMC1, TMEM242), highlighting its potential for biological mechanism elucidation and prognostic assessment. The source code is available at https://github.com/Liangyushi/LaCONIC.

## I. Introduction

CANCER, as a leading cause of mortality worldwide, is characterized by pronounced biological heterogeneity, and its initiation and progression are driven by perturbations in multi-layered molecular regulatory networks [1]–[3]. In this context, constructing accurate survival prediction models is crucial for risk assessment, prognostic stratification, and individualized treatment.

Early cancer survival prediction models, such as Cox proportional hazards model (CoxPH) [4], LASSO-Cox [5] and Coxnet [6] primarily relied on clinical characteristics (e.g., age, sex, subtype, stage) and classical statistical techniques. These methods offered good interpretability, but depended on manual feature construction and linear assumptions, limiting their ability to capture higher-order nonlinear patterns. With the advent of neural networks, deep survival models such as DeepSurv and DeepHit were proposed. DeepSurv replaced the linear risk function with a multilayer neural network to learn complex risk representations [7], while DeepHit discretized continuous time to directly model the joint distribution of survival time and competing risks [8]. Subsequently, Transformer-based methods (e.g., SurvTRACE [9], SSTrans-STG [10]) further exploited the capacity of Transformers to capture sequences and long-range temporal dependencies [11], thereby improving performance in long-term follow-up and competing-risk settings. However, these approaches relied on manually specified time intervals and remained limited in their ability to capture tumor molecular heterogeneity.

With the rapid development of high-throughput sequencing technologies, the biological landscape of cancer has been systematically profiled across multiple molecular layers, including the genome, transcriptome, epigenome, and proteome [12], [13]. Accumulating evidence has revealed pronounced molecular heterogeneity in cancer and demonstrated that such heterogeneity profoundly influences clinical outcomes [14]– [16]. This indicated that clinical information or single-omics data alone are insufficient to capture the key molecular characteristics required for clinical decision-making. Meanwhile, large public resources such as The Cancer Genome Atlas (TCGA) [17] and the International Cancer Genome Consortium (ICGC) [18] have provided rich multi-omics datasets across diverse cancer types, enabling extensive multilayer molecular studies. On this basis, numerous multi-omics survival models have been proposed, such as DCAP [19], HFBSurv [20], CAMR [21], and PCLSurv [22]. Most of these methods were built under a continuous-time framework and achieved improved predictive performance and better characterization of cancer molecular heterogeneity compared with clinic feature-based approaches. However, they generally fail to adequately incorporate the topological characteristics of gene regulatory networks and the impact of cancer subtypes on survival heterogeneity, thereby limiting predictive accuracy and biological interpretability.

In recent years, numerous studies have shown that dysregulation of competing endogenous RNA (ceRNA) networks disrupt miRNA-mediated post-transcriptional regulatory homeostasis, leading to aberrant oncogene activation and tumor suppressor gene silencing, and thereby promoting tumor initiation, progression, and metastasis [23]–[25]. Moreover, different tumor subtypes often exhibit subtype-specific miRNA–ceRNA regulatory modules that are closely associated with patient prognosis, subtype identity, and therapeutic sensitivity [26]– [28]. These observations motivate the joint incorporation of ceRNA network information and cancer subtype annotations into multi-omics survival models, both to more faithfully integrate the regulatory roles of noncoding RNAs and to explicitly preserve subtype-related structural differences during representation learning, thereby enhancing model generalizability and biological consistency.

In this study, we propose LaCONIC, a graph-guided and label-aware multi-omics collaborative learning framework for cancer survival prediction. This framework systematically learn how molecular regulatory networks and cancer subtypes jointly relate to clinical outcomes from a multi-omics perspective, thereby enabling multi-level information integration, survival risk prediction, and biological interpretability. Specifically, a gene regulatory network integrating multiple molecular types and diverse relationships is first constructed to comprehensively characterize complex inter-molecular regulatory patterns. Based on this network, a heterogeneous graph self-supervised pretraining module is developed, which employs node-masking prediction and contrastive learning to generate unified graph-based gene representations that capture both biological and structural information. An adaptive cross-omics representation learning module is then presented, which employs multi-head attention and gating mechanisms to integrate the learned graph-based gene embedding with gene expression data. This module fuses multiple modalities within each omics and performs cross-omics modeling through coordinated components, yielding biologically interpretable latent representations. To explore higher-order dependencies among moleculars, a graph-guided cross-omics interactive learning module is designed to model gene interactions across omics using a Transformer-based structure. Finally, a label-aware constraint mechanism is introduced, integrating subtype classification with multi-level contrastive learning to enhance survival prediction while improving the discriminability and consistency of latent features. Experimental results on multiple cancer datasets demonstrate that LaCONIC outperforms existing methods in various metrics such as Harrell’s CIndex. Interpretability analysis combining SHAP and univariate CoxPH on breast cancer identified key survival-associated regulatory modules and biomarkers, providing insights into cancer heterogeneity and targeted therapy.

## II. Related work

Cancer survival prediction models aim to estimate the survival probability or recurrence risk of patients based on clinical and multi-omics characteristics, thereby providing a quantitative reference for clinical decision-making. Next, we introduce existing methods from the following three aspects:

### A. Based on clinic features

Classical cancer survival prediction models are predominantly built on clinical and pathological features, with the semi-parametric Cox proportional hazards model (CoxPH) as a representative example [4]. On this basis, LASSO-Cox [5] and ElasticNet-Cox (Coxnet) [6] were developed to achieve variable selection and stable modeling via L1/L2 regularization, while Random Survival Forests (RSF) was proposed to capture nonlinear effects and high-order interactions and to handle censored and missing data [29]. With the advancement of deep learning, DeepSurv replaced the linear risk function with a neural network to capture more complex covariate–risk relationships [7], while DeepHit relaxed the proportional hazards assumption by discretizing time and directly modeling the joint distribution of survival time and competing risks [8]. Based on DeepHit, Transformer-based survival models have been proposed to capture global temporal dependencies [30], while SurvTRACE extends this work to model multiple competing events [9], and subsequent semisupervised frameworks SSTrans-STG improve generalization in small-sample and highly censored settings [10]. Although these advanced transformer-based methods perform well in modeling temporal dynamics, they rely on manually defined time bins whose resolution directly affects performance and interpretability. Moreover, their exclusive reliance on clinical variables makes it difficult to capture the molecular heterogeneity of tumors.

### B. Based on multi-omics features

With advances in high-throughput sequencing and integrative multi-omics profiling, survival models have shifted from limited clinical or single-omics features to integrating multiomics data to better capture tumor molecular heterogeneity. For example, Cox-nnet structurally integrates neural networks and regularization mechanisms, enhancing the ability of multiomics modeling [31]. DCAP introduces an autoencoder structure to achieve multi-modal feature dimensionality reduction and reconstruction [19]. TransSurv uses a Transformer to fuse histopathology images and genomic features [32]. HFBSurv adopts a bilinear factorization for multi-omics data integration to capture higher-order interactions [20]. FGCNSurv combines factorized bilinear models with graph convolutional networks to enhance prediction accuracy [33]. CAMR introduces a cross-aligned multi-omics representation learning network that generates modal invariant representations to improve robustness [21]. PCLSurv adopts prototype-based contrastive learning to capture high-level semantic relations that effectively distinguishs samples [22]. Although these models can effectively improve the performance, they lack explicit modeling of gene regulatory relationships and their biological interpretability remains limited.

### C. GRN-informed

With an increasingly refined understanding of tumor molecular mechanisms, there is a growing demand to explicitly encode gene regulatory structures as prior knowledge in survival prediction models, thereby enhancing discriminative performance while providing traceable biological interpretations. For example, Sun et al. introduced a graph Laplacian penalty into the Cox framework to smooth the coefficients of neighboring genes [34]. Liu et al. inferred sample-level pathway activities based on pathway topology to strengthen mechanistic relevance [35]. Within deep learning–based survival frameworks, KEGG pathway structures have been encoded as intermediate representations to enable end-to-end interpretable learning [36], while GRN priors have been integrated into multi-omics training to identify key regulatory nodes [37]. Although these approaches help improve robustness and biological interpretability at the gene level, the utilization of post-transcriptional regulatory networks (e.g., ceRNA) for risk stratification remains limited.

Overall, these methods represent a shift in survival prediction from feature-driven to network-driven modeling, moving from single clinical data to multi-omics integration and structured modeling. Nevertheless, existing models remain insufficient in jointly characterizing multi-omics features, gene regulatory network structures, and cancer subtypes. This makes them difficult to fully capture the molecular heterogeneity of cancer and limit their predictive performance and biological interpretability. Thus, a unified framework that jointly integrates these components to model complex molecular interactions is critically needed.

## III. Materials and methods

The LaCONIC collaboratively integrates multi-omics expression data, gene regulatory network structure, and cancer subtype into a unified deep survival prediction framework. The overall framework of LaCONIC is shown in Figure 1.

**Fig. 1.**
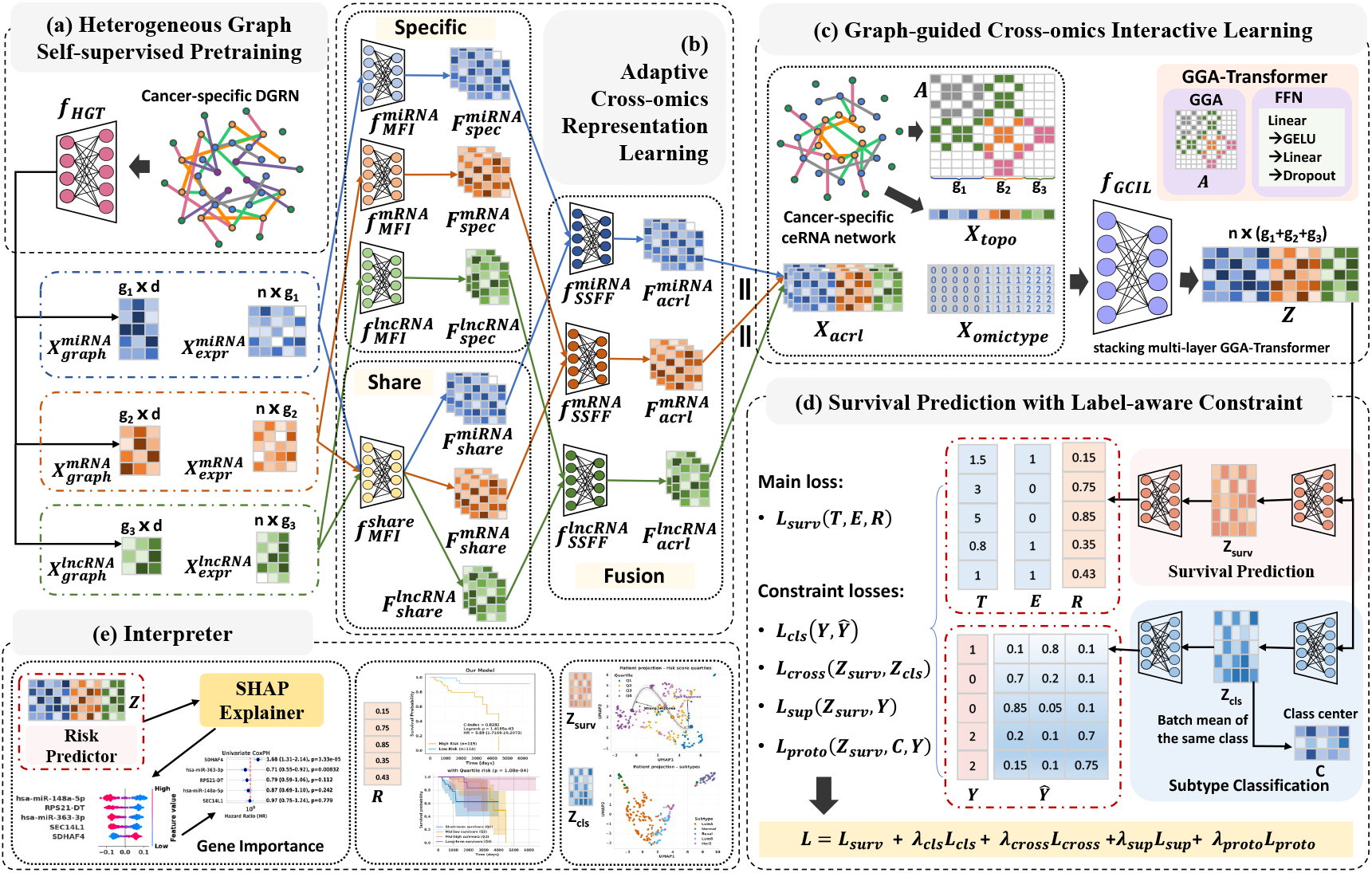
Overview of the LaCONIC framework. (a) Heterogeneous Graph Self-supervised Pretraining: an HGT-based network learns molecular representations that integrate biological semantics and topological structure. (b) Adaptive Cross-omics Representation Learning: multi-head attention and gating collaboratively model omics-specific and shared features to adaptively align and fuse multi-omics information, yielding robust and biologically interpretable latent representations. (c) Graph-guided Cross-omics Interaction Learning: a graph-guided Transformer models cross-omics gene–gene interactions, capturing crosslayer dependencies and reflecting molecular heterogeneity across patients. (d) Survival Prediction with Label-aware Constraint: survival risk prediction is jointly optimized with subtype classification and multi-level contrastive learning to enhance the discriminability and consistency of the latent feature space. (e) Interpreter: SHAP and univariate Cox regression are used to identify survival-associated key genes and support downstream analyses such as risk stratification.

### A. Dataset

We first systematically downloaded and curated multisource relational data for miRNAs, mRNAs, lncRNAs, and disease entities (see Supplementary Table S1 for details), and constructed a large-scale disease gene regulatory network (DGRN) containing seven molecular associations: miRNA-mRNA, miRNA-lncRNA, mRNA-mRNA, mRNAdisease, miRNA-disease, lncRNA-disease, and miRNAmiRNA. Among these, the miRNA-miRNA associations were inferred following the previous research [38], [39]. By comprehensively considering the regulatory effects of common target genes and the influence of protein-protein interaction patterns on miRNA synergy, the high-confidence functional synergistic relationships between miRNAs were constructed.

Then the R package TCGAbiolinks was adopted to obtain survival data and miRNA/mRNA expression profiles for 11 cancer types from TCGA. Based on subtype label overlap, we constructed 5 single-cancer and 2 pan-cancer datasets (Table I). From the large DGRN resources, we further extracted cancer-specific DGRN and ceRNA networks to capture differences in regulatory structure across cancers (network statistics are in Supplementary Table S2). To focus on cancer-relevant biomarkers, we adopted disease-specific survival (DSS) events and times as the final survival labels instead of overall survival (OS). High-confidence subtype labels were derived by intersecting three authoritative sources: (a) subtypes defined in [40]; (b) clinical subtype data from the TCGA Pan-Cancer Atlas (PanCanAtlas) via cBioPortal [41]; and (c) molecular subtype annotations from PanCanAtlas on the UCSC Xena platform [42]. This multi-source integration yielded a unified multi-cancer dataset comprising expression profiles, clinical information, and regulatory networks. More details on data preprocessing are in Supplementary Material Section 1.

**TABLE 1.** Statistical overview of multi-omics, ceRNA network and clinical data across cancer types.

| Cancer | Abbr. |  | Sample |  | miRNA | mRNA | lncRNA | Event | Censored | Label | ceRNA |
| --- | --- | --- | --- | --- | --- | --- | --- | --- | --- | --- | --- |
| Breast invasive carcinoma | BRCA |  | 1143 |  | 116 | 4129 | 141 | 103 | 1040 | 5 | 561,289 |
| Cervical squamous cell carcinoma and endocervical adenocarcinoma | CESC |  | 255 |  | 78 | 1849 | 120 | 47 | 208 | 2 | 127,028 |
| Head and Neck squamous cell carcinoma | HNSC |  | 461 |  | 118 | 1537 | 55 | 125 | 336 | 2 | 95,099 |
| Sarcoma | SARC |  | 224 |  | 174 | 2400 | 90 | 69 | 155 | 4 | 286,299 |
| Uterine Corpus Endometrial Carcinoma | UCEC |  | 494 |  | 70 | 2931 | 165 | 54 | 440 | 4 | 259,070 |
| Colon adenocarcinoma | COAD | COAD_ESCA_READ_STAD | 312 | 921 | 69 | 1554 | 43 | 179 | 742 | 6 | 91,982 |
| Esophageal carcinoma | ESCA |  | 167 |  |  |  |  |  |  |  |  |
| Rectum adenocarcinoma | READ |  | 107 |  |  |  |  |  |  |  |  |
| Stomach adenocarcinoma | STAD |  | 335 |  |  |  |  |  |  |  |  |
| Glioblastoma multiforme | GBM | GBM_LGG | 142 | 638 | 193 | 6285 | 376 | 216 | 422 | 3 | 1,280,772 |
| Brain Lower Grade Glioma | LGG |  | 496 |  |  |  |  |  |  |  |  |

### B. Heterogeneous Graph Self-supervised Pretraining

To fully capture the complex semantic relationships among miRNAs, mRNAs, lncRNAs, and diseases, we design a heterogeneous graph self-supervised pretraining module based on the Heterogeneous Graph Transformer (HGT) [43]. This module jointly optimizes masked node prediction and contrastive learning to learn unified node embeddings across omics and relation types in an unlabeled setting, providing high-quality network feature initialization. Concretely, given a heterogeneous graph 𝒢 = (𝒱, *ℰ*) with multiple node and edge types, we initialize type-specific embeddings *E*^*t*^HGT for each node type *t* ∈ {miRNA, mRNA, lncRNA, disease} using an independent embedding network Embedding^*t*^HGT, and feed them into a heterogeneous graph encoder Encoder composed of two HGTConv layers to obtain the final node representations *H* = *f*_HGT_(𝒱, ℰ).

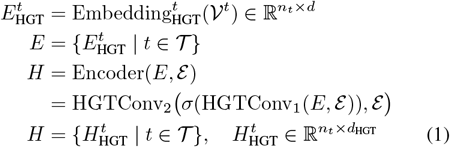

where *n*_*t*_ denotes the number of nodes of type *t, d* is the initial node feature dimension, and *d*_HGT_ is the output embedding dimension. The function *σ*(·) denotes the GELU activation. HGTConv is the core operator of HGT, which captures crosstype interactions via meta-relation–specific attention. The output *H* contains the embeddings of all node types.

To encourage LaCONIC to learn local context-aware semantics, we randomly mask a proportion *r* of nodes for each node type (*r* = 0.15) and require the model to reconstruct their original indices from the graph context. For each node type *t*, we define an index decode 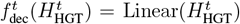. A cross-entropy loss over masked nodes is then used as the masked node prediction objective.

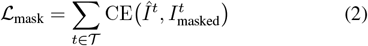

where 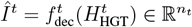 is the reconstructed probability distribution over node indices of type *t*, and 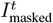 denotes the ground-truth index labels for the masked nodes.

To enforce global consistency of node embeddings across different views, we adopt a SimCLR [44] contrastive learning objective. Specifically, we generate two randomly masked views for each node and obtain the corresponding representations 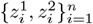. The contrastive loss is defined as:

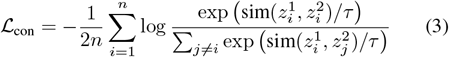

where sim(*a, b*) = *a*^⊤^*b* denotes the dot-product similarity and *τ* is a temperature parameter. In this task, all node types are jointly included in the contrastive set, i.e., *n* denotes the total number of nodes across all types, which encourages global consistency modeling across different biological entity types.

We then jointly optimize these two self-supervised objectives to obtain the overall pretraining loss:

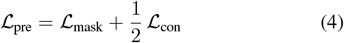

Through this pretraining strategy, we obtain the high-quality initial molecular graph features 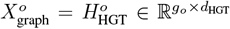 where *g*_*o*_ denotes the number of genes (molecular entities) in omics *o* ∈ {miRNA, mRNA, lncRNA}.

### C. Adaptive Cross-omics Representation Learning

To fully exploit the complementarity and consistency of multi-omics, multi-modal data, LaCONIC introduces an Adaptive Cross-omics Representation Learning (ACRL) module that performs intra-omics multi-modal fusion and interomics alignment to obtain robust, biologically interpretable latent representations. ACRL comprises three components: an omics-specific encoder, an omics-shared encoder, and a specific–shared fusion layer. Both encoders share a Multiview Feature Integration (MFI) backbone: the omics-specific encoder uses separate MFI networks for each omics to capture modality-specific patterns, whereas the shared encoder uses a common MFI network across omics to align their latent spaces and learn cross-modal commonalities. The fusion layer then employs a gating mechanism to combine specific and shared features into representations that preserve omics specificity while enforcing cross-omics consistency.

#### 1) Omics-specific encoder

The core component of omics-specific encoder is the Multi-view Feature Integration (MFI) network, whose key idea is to jointly learn intra-omic multimodal representations from gene expression and graph-based features via a gating mechanism and multi-head attention. On this basis, it instantiates an independent, parameterized MFI network 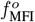 for each omics *o*, ensuring that LaCONIC can learn omics-specific feature representation for each omic. Specifically, for the *o*-th omics, the inputs consist of a gene expression matrix 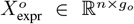 and a gene-level graph feature matrix 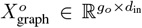, where *n* is the number of samples and *d*_in_ = *d*_HGT_ denotes the initial graph feature dimension. The MFI module 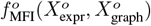 is composed of the following four components:

##### Intra-modal: feature projection

For each omics *o*, we first replicate the precomputed graph-based gene representations across all samples, yielding 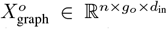. Similarly, the expression matrix is broadcast along the gene dimension to 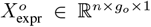 and then passed through a gene-wise mapping layer 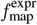 to project it into the same feature space as the graph representation 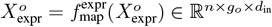.

The resulting expression features 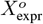 are then fed into an expression-specific projection layer 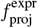 and a shared projection layer 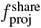 to obtain expression-specific representations 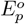 and shared representations 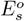, seperately:

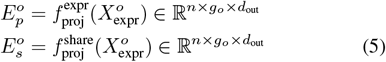

Analogously, the replicated graph features 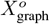 cessed by a graph-specific projection layer 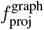 are proand the same shared projection layer 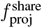, yielding graph-specific and shared graph representations as:

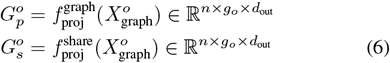

where 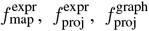 and 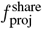 are all implemented as single-layer blocks consisting of a linear transformation, Layer Normalization (LN), and a GELU activation.

##### Inter-modal: shared features fusion

To balance the contributions of expression and graph features to the shared representation, 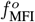 first concatenates the two shared components, and feeds 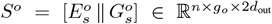 into a channelwise gating layer 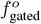 to achieve adaptive fusion: To balance the contributions of expression and graph features to the shared representation, 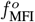 first concatenates the two shared components to form 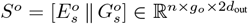, and then feeds *S*^*o*^ into a channel-wise gating layer 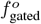 to achieve adaptive fusion as:

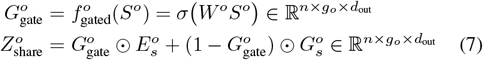

where *σ*(·) denotes the sigmoid function, 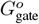 is the learned gating weight, and ⊙ denotes element-wise multiplication.

##### Intra-inter modal: multi-modal feature fusion

We first combine the shared representation with the modality-specific components and normalize the results to obtain enhanced expression and graph features. These are then concatenated to form a unified multi-view representation *H*^*o*^ as:

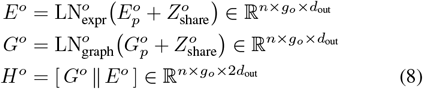

##### Intra-omics: gene interaction learning

To capture higherorder gene dependencies within each omics, 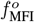 applies multi-head self-attention MHA to *H*^*o*^, followed by residual connection and normalization, yielding the final omics-specific integrated representation 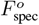 for omic *o*:

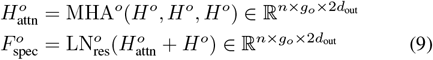

#### 2) Omics-shared encoder

This encoder is designed to align latent representation distributions across different omics, thereby capturing cross-modal consistency. Similar to the omics-specific layer, it is also built upon the MFI architecture, but adopts a parameter-sharing strategy. In other words, all omics share the same MFI network 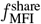, which enforces distributional alignment and semantic consistency in a common feature space. Formally, the shared representations for *o* omic can be written as:

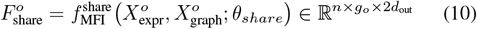

where *θ*_*share*_ is a shared parameter for all omics.

#### 3) Specific-shared fusion layer

This layer is designed to integrate representations from the omics-specific and omics-shared encoders, yielding final features that jointly preserve modality-specific information and cross-omics consistency. Its core component is the Specific-Shared Feature Fusion (SSFF) module. SSFF first performs channel-wise gated fusion between specific and shared representations, and then applies Dropout and Layer Normalization (LN) to obtain a stable and fused omics-level representation. Formally, given the omics-specific and shared features for the *o*-th omics, the fusion process implemented by 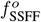 is defined as:

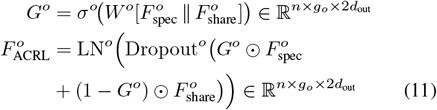

This mechanism adaptively modulates the relative contributions of specific and shared information for each feature channel, enabling fine-grained information balancing.

### D. Graph-guided Cross-omics Interactive Learning

The Graph-guided Cross-omics Interactive Learning (GCIL) module aims to learn context-dependent and cross-modal gene interactions under the guidance of the ceRNA network, enabling the model to capture key regulatory genes and their global impact on cancer prognosis. Before feeding data into GCIL, we first concatenate miRNA, mRNA, and lncRNA features along the gene dimension to obtain 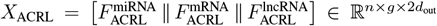 where *g* = *g*_miRNA_ + *g*_mRNA_ + *g*_lncRNA_ denotes the total number of genes. Each gene is additionally assigned an omics-type label *T* = [*t*_miRNA_, *t*_mRNA_, *t*_lncRNA_], *t*_*o*_ ∈ { 0, 1, 2 }, which is used to construct the omics-type feature embedding *X*_omictype_ ∈ ℝ^*n×g*^. Based on the ceRNA network adjacency matrix *A* ∈ ℝ^*g×g*^, we further compute the betweenness centrality of each gene to obtain a topological feature vector *X*_topo_ ∈ ℝ^*g×*1^ (Specific calculation process can be found in Supplementary Material Section 2). Thus, given *X*_ACRL_, *X*_omictype_, *X*_topo_, and *A*, the GCIL module produces sample-level representations as:

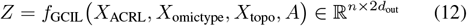

In the following, we detail the architecture of *f*_GCIL_.

#### 1) Graph-guided attention mechanism

To explicitly model regulatory dependencies among genes, GCIL introduces a Graph-Guided Attention (GGA) mechanism, which augments standard multi-head self-attention with structural priors from the ceRNA adjacency matrix *A*. Formally, given input features 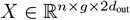 and adjacency matrix *A* ∈ ℝ^*g×g*^, the attention weights are defined as:

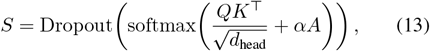

where *Q* = reshape(*W*_*q*_*X*), *K* = reshape 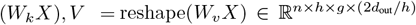 are the query, key, and value tensors for *h* attention heads, 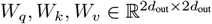 are learnable projection matrices, *d*_head_ = 2*d*_out_*/h* is the per-head dimensionality, and *α* is a learnable bias coefficient that controls the strength of the graph structural prior. GGA can guide models to focus more on known biological regulatory connections while retaining the ability to explore new potential connections. Then, the graph-guided contextual features are obtained by aggregating the value tensor with the attention weights, followed by a linear projection as:

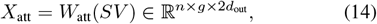

where 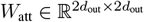 is a learnable parameter matrix.

#### 2) GGA-Transformer backbone encoder

By integrating GGA with Transformer, GCIL constructs a GGA-Transformer backbone consisting of a GGA mechanism and a two-layer feed-forward network (FFN) for nonlinear feature transformation. GCIL models cross-omics gene interactions by stacking multiple GGA-Transformer encoder layers, with residual connections and layer normalization applied in each layer to stabilize training and enhance representational capacity.

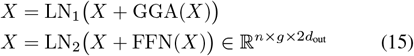

To capture semantic differences arising from omics type and topological context, we further introduce two types of embeddings:

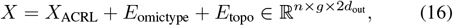

where 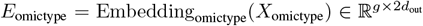 is the omics-type embedding, and 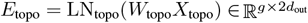 is the linearly projected and normalized topological embedding, with *W*_topo_ being a learnable weight matrix.

Finally, GCIL applies a gated pooling scheme that combines average and max pooling across the gene dimension to obtain sample-level representations as:

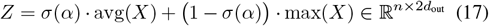

where *α* is a learnable scalar, *σ*(·) is the sigmoid function, and avg(·) and max(·) denote average and max pooling over the gene dimension, respectively.

### E. Survival Prediction with Label-aware Constraint

LaCONIC jointly optimizes survival prediction and label-aware constraint to maintain high survival prediction accuracy while learning structurally coherent and biologically interpretable multi-omics representations.

#### 1) Survival prediction

This constitutes the core optimization objective of LaCONIC and is used to learn continuous representations that reflect patient-specific prognostic risk. Given the fused multi-omics representation *Z*, a survival head *f*_surv_ based on multilayer perceptron (MLP) is employed to map *Z* nonlinearly to scalar risk scores *R* = *f*_surv_(*Z*) = {*r*_1_, …, *r*_*i*_, …, *r*_*n*_ }, where *r*_*i*_ is the risk score of the i-th sample. To balance the contributions of events and censored observations, we design a weighted Cox loss as:

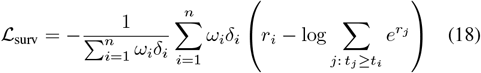

where *δ*_*i*_ ∈ {0, 1} is the event indicator (*δ*_*i*_ = 1 if the event occurs), *t*_*i*_ is the observed survival time, and *ω*_*i*_ is the sample weight. In practice, we set 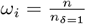 for event samples (*δ*_*i*_ = 1) and *ω*_*i*_ = 1 otherwise. This weighting scheme upweights rare event cases, increasing the model’s sensitivity to failure events and stabilizing gradient estimation during training.

#### 2) Label-aware constraint

This constraint includes a subtype classification loss and three contrastive objectives, which encourages both consistency and discriminability of the learned representations, thereby imposing explicit regularization on the feature space and improving survival prediction.

##### Subtype classification constraint

The cancer subtype classification task acts as a supervised auxiliary signal that encourages biologically coherent and well-separated subtype representations in the latent space, providing complementary supervision for survival risk modeling. An MLP-based classifier *f*_cls_ maps the fused representation *Z* to subtype logits *Ŷ* = *f*_cls_(*Z*) ∈ ℝ^*n×K*^, where *K* is the number of subtypes. To emphasize hard examples and alleviate class imbalance, we adopt the Focal Loss [45]:

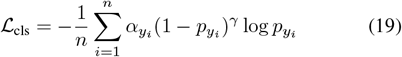

where *y*_*i*_ is the ground-truth of sample 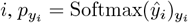 is the predicted probability of the true class, and *α* = 1, *γ* = 2.0 are the class-balancing and focusing parameters.

##### Multi-level contrastive constraint

To further enforce the consistency between survival representations and subtype features, this constraint introduces three following contrastive losses to jointly enhance the global consistency and local discriminability of the representation space.

· Cross-task feature alignment loss. This term aligns the survival and classification representations. Given two task-specific embeddings *Z*_surv_ = *z*_surv,*i*_ and *Z*_cls_ = {*z*_cls,*i*_} for each sample, we maximize their mutual information via a bidirectional InfoNCE loss [46] as:

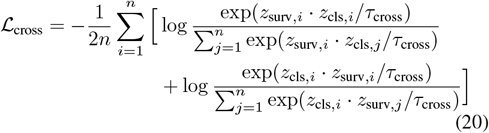

where *τ*_cross_ denotes the temperature, and *Z*_surv_ and *Z*_cls_ are the *ℓ*_2_-normalized penultimate-layer embeddings of the survival head *f*_surv_ and classification head *f*_cls_, respectively.

· Label-supervised contrastive loss. This loss imposes a label-informed discriminative structure on the survival embedding space by pulling same-subtype samples together and pushing different subtypes apart. For each anchor *i*, given normalized survival embeddings *z*_surv,*i*_, ground-truth labels *y*_*i*_, and the positive set *P* (*i*) = {*p* ≠ *i* |*y*_*p*_ = *y*_*i*_ }, the supervised contrastive loss [47] is defined as:

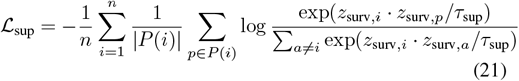

· Prototype-based contrastive loss. This loss is adopted to enhance the stability of decision boundaries and mitigate cross-batch drift. For each subtype *k*, LaCONIC maintains a prototype vector *c*_*k*_ in the classification embedding space, initialized from the output of *f*_cls_ and updated via Exponential Moving Average (EMA) [48]. Let *B*_*k*_ = *i* | *y*_*i*_ = *k* be the set of samples of subtype *k* in the current mini-batch, their mean embedding is defined as:

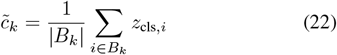

Then the global prototype is updated by 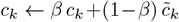 with momentum *β* ∈ [0, 1).

Given a survival embedding *z*surv, *i* and its class prototype 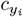 as the positive (all other prototypes as negatives), the prototype-based contrastive loss [49] is defined as:

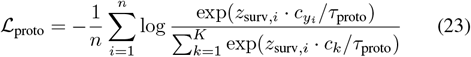

where *τ*_proto_ is a temperature parameter and *K* is the number of subtypes. By combining the above loss terms, the overall objective of LaCONIC is defined as:

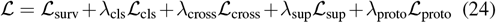

where *λ*_cls_, *λ*_cross_, *λ*_sup_, and *λ*_proto_ are the weighting coefficients for the corresponding auxiliary objectives.

### F. SHAP-based Gene Importance

To enhance model interpretability, we employ SHAP (SHapley Additive exPlanations) [50] to quantify the contribution of each gene feature to survival prediction. SHAP is a posthoc, game-theoretic explanation method that computes Shapley values by estimating the average marginal contribution of each feature over all possible feature coalitions, thereby revealing how individual features influence the model output. Concretely, we initialize a `shap.GradientExplainer` using the trained survival head *f*_surv_ together with the training representations 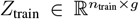. We then feed the test representations 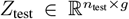 into the explainer to obtain a SHAP value matrix 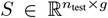, where *S*_*ij*_ denotes the marginal contribution of gene *j* to the predicted risk of sample Based on *S*, we compute the mean absolute SHAP value for each gene across all test samples, use this quantity as the gene importance to rank the genes, and then select the top 20 genes for downstream biological validation to investigate their potential regulatory roles in cancer prognosis.

## IV. Experiments

### A. Experimental setup

We systematically evaluated LaCONIC using multiple complementary survival analysis metrics, including Harrell’s concordance index (Harrell’s C-Index) to assess ranking concordance of individual survival times, the time-dependent area under the ROC curve integrated over time (iAUC) to quantify discriminative ability at the 25th, 50th, and 75th percentiles of the survival time distribution, the inverse probability of censoring weighting C-Index (IPCW C-Index) to reduce bias from censoring, and Kaplan–Meier (KM) survival curve analysis with median risk–based high/low risk stratification, where group differences were assessed by the log-rank test. To ensure representative and stable multimodal data, we adopted stratified splitting based on joint strata of event status and subtype label, first dividing each dataset into 80% training and 20% test, and then holding out 20% of the training set for validation (64%/16%/20% train/validation/test). More detailed experimental settings are provided in the Supplementary Material Section 3.

### B. Evaluation of survival prediction performance

To evaluate the performance of LaCONIC, we compared it against 14 existing methods, including three classical survival models (Coxnet, CoxBoost [51], RSF), four deep discrete-time models (DeepHit, Transformer Survival, Trans-STG, SurvTRACE), and seven deep continuous-time models (DeepSurv, Cox-nnet, DCAP, HFBSurv, CAMR, FGCNSurv, PCLSurv). To ensure a fair comparison, all methods except FGCNSurv and PCLSurv (which were inherently designed to use only miRNA and mRNA) were trained on the same multi-omics inputs. Moreover, except for HFBSurv, CAMR, FGCNSurv, and PCLSurv (which employed a specific encoder for each omics), the remaining baselines took the concatenated three-omics gene expression matrix as input.

#### 1) Survival ranking concordance

We evaluated all methods on seven cancer datasets (five single-cancer and two pan-cancer) and reported their Harrell’s C-Index values in Table II, which depicted that LaCONIC achieves the highest C-Index on all datasets except UCEC, with particularly pronounced gains on CESC (0.8943), SARC (0.8844), and the two pan-cancer datasets COAD ESCA READ STAD (0.7603) and GBM LGG (0.8893), clearly surpassing all baselines. Traditional methods performed markedly worse, especially on complex multi-cancer data. For example, CoxNet attained only a C-Index of 0.5242 on COAD ESCA READ STAD, far below LaCONIC. Discrete-time models improved over traditional approaches but still fall short of LaCONIC. For instance, Trans-STG obtained C-Index values of 0.7940 on SARC and 0.8377 on CESC, compared with 0.8844 and 0.8943 for LaCONIC. Continuous-time models were competitive on some datasets but do not outperform LaCONIC overall. For example, PCLSurv reached 0.8219 on BRCA, close to LaCONIC, yet remained inferior on CESC and GBM LGG (0.7434 and 0.8683 vs. 0.8943 and 0.8893).These results demonstrated that LaCONIC provides consistently superior survival ranking concordance across both single-cancer and multi-cancer settings.

**TABLE 2.** Comparison of different cancer survival prediction models based on Harrell’S C-Index across seven cancer datasets. Boldface and underlining indicate the best and second-best performances, respectively.

|  | Model | BRCA | CESC | HNSC | SARC | UCEC | COAD_ESCA_READ_STAD | GBM_LGG | Mean±STD |
| --- | --- | --- | --- | --- | --- | --- | --- | --- | --- |
| Traditional | Coxnet | 0.8120 | 0.7698 | 0.6117 | 0.8116 | 0.6914 | 0.5242 | 0.6455 | 0.6952±0.11 |
|  | CoxBoost | 0.7549 | 0.7415 | 0.6146 | 0.7387 | 0.7360 | 0.5991 | 0.8385 | 0.7176±0.08 |
|  | RSF | 0.7584 | 0.7245 | 0.5836 | 0.7814 | 0.7343 | 0.6784 | 0.8280 | 0.7269±0.08 |
| Based on discrete time | DeepHit | 0.6860 | 0.7132 | 0.6830 | 0.7487 | 0.7145 | 0.6921 | 0.6876 | 0.7036±0.02 |
|  | Transformer Survival | 0.7542 | 0.8566 | 0.6787 | 0.8065 | <u>0.8399</u> | 0.6905 | 0.7443 | 0.7673±0.07 |
|  | Trans-STG | 0.8002 | 0.8377 | <u>0.7061</u> | 0.7940 | **0.8449** | 0.7176 | <u>0.8799</u> | 0.7972±0.07 |
|  | SurvTRACE | 0.8071 | <u>0.8679</u> | 0.6499 | 0.8367 | **0.8449** | 0.7158 | <u>0.8606</u> | <u>0.7975±0.08</u> |
| Based on continuous time | DeepSurv | 0.6762 | 0.7887 | 0.6477 | 0.4899 | 0.7079 | 0.5718 | 0.6417 | 0.6463±0.10 |
|  | Cox-nnet | 0.7402 | 0.6868 | 0.4820 | 0.7889 | 0.6766 | 0.6604 | 0.7587 | 0.6848±0.10 |
|  | DCAP | **0.8282** | 0.8038 | 0.6398 | <u>0.8593</u> | 0.7162 | 0.7222 | 0.8203 | 0.7700±0.08 |
|  | HFBSurv | 0.7554 | 0.8075 | 0.6477 | <u>0.8116</u> | 0.6865 | 0.6692 | 0.8326 | 0.7443±0.08 |
|  | CAMR | 0.8041 | 0.7642 | 0.6001 | 0.8178 | 0.7186 | 0.7011 | 0.8420 | 0.7497±0.08 |
|  | FGCNSurv | 0.8091 | 0.8264 | 0.6037 | 0.8166 | 0.6898 | 0.6977 | 0.8655 | 0.7584±0.10 |
|  | PCLSurv | <u>0.8219</u> | 0.7434 | 0.6441 | 0.8116 | 0.7393 | <u>0.7263</u> | 0.8683 | 0.7650±0.07 |
|  | LaCONIC (our) | **0.8282** | **0.8943** | **0.7075** | **0.8844** | 0.8218 | **0.7603** | **0.8893** | **0.8266±0.07** |

Additionally, Supplementary Figure S1 summarized iAUC and IPCW C-Index distributions for all models across seven datasets, ordered by decreasing mean performance. LaCONIC achieved the highest mean *iAUC* (0.8563 ± 0.07), exceeding the second-best method SurvTRACE (0.8220 ± 0.09) by 3.43%. In terms of *IPCW C-Index*, LaCONIC (0.7766 *±* 0.08) again exhibited the leading performance, improving on Trans-STG (0.7188 *±* 0.11) by 5.78% and substantially outperforming conventional methods. These findings indicated that LaCONIC attained high discriminative accuracy and robustness across diverse cancer survival prediction tasks.

#### 2) Kaplan-Meier risk stratification

To further assess risk stratification ability, we performed KM survival analysis for all models on seven cancer datasets. Table III reported the corresponding log-rank test p-values, where smaller values indicated more pronounced separation between risk groups. LaCONIC achieved statistically significant risk stratification (p*<*0.05) on all datasets, with the smallest p-values observed on CESC (p=1.27E-03), HNSC (p=3.98E-02) and SARC (p=1.62E-04), and its mean p-value (0.0063 *±* 0.01) was markedly lower than that of any baselines. In contrast, traditional models and other deep approaches often yielded relatively large p-values on several datasets (e.g., HNSC and UCEC), indicating limited risk stratification capability. Supplementary Figure S2 further illustrated the KM survival curves of all models on the SARC dataset. LaCONIC produced the clearest separation between high and low risk groups, with the high-risk group’s survival dropping rapidly and an HR of 17.81, indicating nearly 18-fold higher mortality risk and demonstrating clinically meaningful risk stratification.

**TABLE 3.** Log-RANK TEST P-VALUES FROM Kaplan–Meier (KM) risk stratification for all models on the seven cancer datasets, where underlining denotes non-significant results and boldface denotes the most significant separation.

|  | Model | BRCA | CESC | HNSC | SARC | UCEC | COAD_ESCA_READ_STAD | GBM_LGG | Mean±STD |
| --- | --- | --- | --- | --- | --- | --- | --- | --- | --- |
| Traditional | Coxnet | 2.98E-04 | 1.00E-02 | <u>1.25E-01</u> | 1.69E-02 | <u>1.64E-01</u> | <u>7.85E-01</u> | 2.54E-02 | <u>0.1610±0.28</u> |
|  | CoxBoost | 3.58E-01 | 2.78E-02 | <u>2.83E-01</u> | 8.02E-03 | 4.30E-02 | 3.65E-02 | 6.18E-10 | <u>0.1080±0.15</u> |
|  | RSF | 4.94E-03 | 5.01E-02 | <u>3.53E-01</u> | 6.33E-02 | 1.43E-01 | 2.78E-05 | 1.34E-10 | <u>0.0877±0.13</u> |
| Based on discrete time | DeepHit | 8.38E-03 | <u>9.97E-02</u> | <u>1.83E-01</u> | <u>2.55E-01</u> | <u>1.03E-01</u> | 9.47E-04 | 7.59E-04 | <u>0.0930±0.10</u> |
|  | Transformer Survival | 3.92E-02 | 2.55E-02 | <u>1.22E-01</u> | <u>9.20E-02</u> | 9.24E-03 | 1.78E-02 | 7.45E-06 | 0.0437±0.05 |
|  | Trans-STG | 6.44E-04 | 2.99E-03 | <u>1.19E-01</u> | 7.99E-03 | 6.32E-03 | 1.67E-05 | 1.26E-11 | 0.0196±0.04 |
|  | SurvTRACE | **1.43E-04** | 1.49E-02 | <u>1.04E-01</u> | 2.07E-03 | **4.30E-04** | 2.77E-04 | **2.50E-13** | 0.0174±0.04 |
| Based on continuous time | DeepSurv | <u>3.06E-01</u> | 2.07E-02 | <u>4.92E-02</u> | <u>7.22E-01</u> | 1.28E-02 | <u>7.18E-02</u> | 3.21E-02 | <u>0.1736±0.26</u> |
|  | Cox-nnet | 6.94E-02 | 3.82E-02 | <u>5.13E-01</u> | 2.55E-03 | 4.66E-01 | 2.30E-03 | 2.00E-06 | <u>0.1560±0.23</u> |
|  | DCAP | 8.48E-04 | 2.28E-02 | <u>1.83E-01</u> | 8.00E-04 | 2.31E-02 | **1.37E-06** | 7.22E-09 | 0.0329±0.07 |
|  | HFBSurv | 2.43E-02 | 7.00E-03 | 4.65E-02 | 2.78E-03 | 6.35E-02 | 2.47E-03 | 9.64E-06 | 0.0209±0.03 |
|  | CAMR | 6.83E-03 | 3.13E-02 | <u>5.21E-01</u> | 1.42E-02 | 2.52E-02 | 3.10E-04 | 2.63E-09 | <u>0.0855±0.19</u> |
|  | FGCNSurv | 4.70E-03 | 4.93E-03 | <u>5.55E-01</u> | 5.15E-03 | 5.40E-02 | 2.57E-03 | 3.45E-12 | <u>0.0894±0.21</u> |
|  | PCLSurv | 4.08E-03 | 1.85E-02 | <u>8.74E-02</u> | 1.89E-03 | <u>1.50E-01</u> | 3.69E-04 | 1.01E-12 | <u>0.0375±0.06</u> |
|  | LaCONIC (ours) | 1.42E-03 | **1.27E-03** | **3.98E-02** | **1.62E-04** | 1.05E-03 | 1.69E-04 | 7.61E-11 | **0.0063±0.01** |

### C. Evaluation of the correlation between clinical covariates and predicted risk

To examine the correlation between LaCONIC-predicted risk and clinical covariates and assess its independent prognostic value, we performed CoxPH regression on each cancer dataset. For each dataset, we assembled survival time, event status, clinical variables (age, sex, stage, subtype, etc.), and the LaCONIC risk score. Continuous variables (e.g., age, risk score) were z-score standardized and categorical variables (e.g., sex, stage) were one-hot encoded. Samples with missing or implausible values were excluded. Univariable and multivariable CoxPH were fitted using the `lifelines` package [52]. For each covariate, we estimated the hazard ratio (HR) and its 95% confidence interval (CI) in the univariable model. Multivariable CoxPH were used to assess whether the LaCONIC risk score preserved prognostic significance after adjustment for clinical factors.

Figures 2 and Supplementary Figure S3 summarized the univariable and multivariable Cox regression results across all cancer datasets. The LaCONIC risk score consistently emerged as a significant and independent prognostic factor. In BRCA, the risk score was strongly associated with survival in both univariable (HR = 3.60, p = 1.59 *×* 10^−4^) and multivariable (HR = 4.49, p = 1.25 *×* 10^−4^) models, whereas clinical variables such as subtype and stage were not significant, indicating that the predicted risk alone captured major survival heterogeneity. Similar patterns were observed in other cancers: in univariable analyses, the HRs for the predicted risk in CESC, SARC, and GBM LGG were 3.70, 2.98, and 5.81, respectively (all p*<*0.05), exceeding those of conventional clinical factors. After adjustment for age, sex, subtype, and other covariates, the LaCONIC risk score remained statistically significant (p*<*0.05), with HRs of 3.91, 7.81, and 6.86 in CESC, SARC, and GBM LGG, respectively, thereby underscoring its role as a key prognostic determinant in survival analysis.

**Fig. 2.**
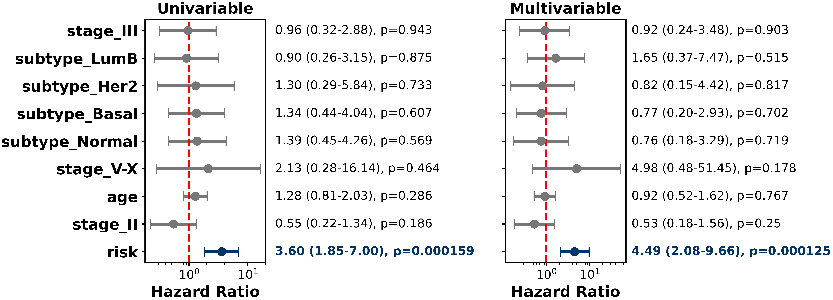
Univariable and multivariable Cox proportional hazards analysis of LaCONIC-predicted risk and clinical covariates on the BRCA dataset. The forest plot displays the hazard ratio (HR), 95% confidence interval (CI), and p-value for each covariate.

### D. Interpretability analysis

To investigate the interpretability of LaCONIC in survival risk prediction, we took the BRCA dataset as an example and conducted a systematic analysis. First, we computed SHAP-based importance to quantify the contribution of each gene to the predicted risk and selected the top 20 genes with the largest impact on the model output (Figure 3a). We then performed univariable CoxPH regression for these genes and identified 14 genes significantly associated with survival (p*<*0.05, Figure 3b). By comparing gene expression levels, SHAP values and the direction of risk in the CoxPH, we found that the correlation between expression and SHAP values was fully consistent with the direction of HR (Figure 3c). Specifically, genes with HR*>*1 (e.g., hsa-miR-365a-3p, TMEM242, ARMC1) showed a positive correlation between expression and SHAP values, indicating that higher expression increased survival risk, whereas genes with HR*<*1 (e.g., hsa-miR-148a-3p, hsa-miR-363-3p, GSTM4) showed a negative correlation, suggesting a protective effect of high expression.

**Fig. 3.**
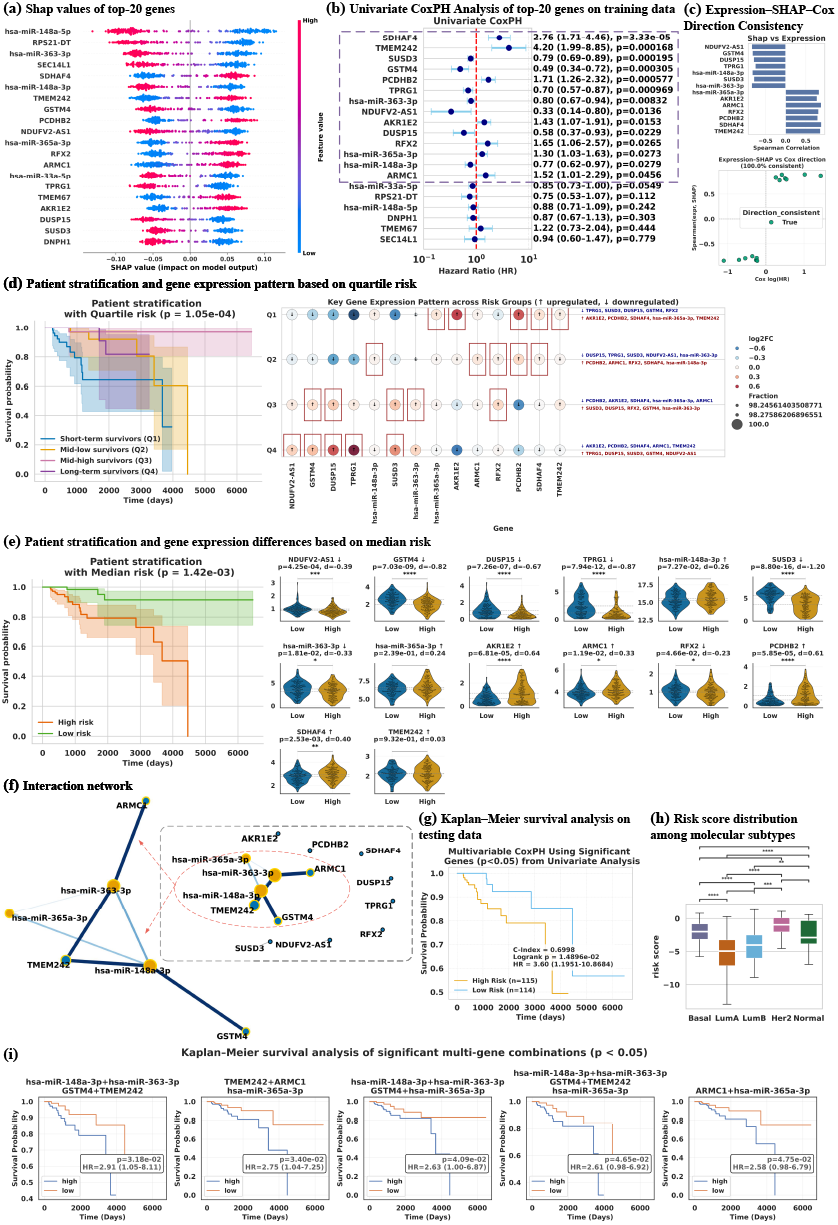
SHAP-based interpretability analysis of the LaCONIC model.

#### 1) Analysis of survival risk and gene expression patterns

We stratified patients into four groups according to the quartiles of the LaCONIC-predicted risk scores: short-term survivors (Q1, *>*75%), mid-low survivors (Q2, 50–75%), mid-high survivors (Q3, 25–50%), and long-term survivors (Q4, *<*25%). We then analyzed survival outcomes and gene expression patterns across these groups. The KM curves in Figure 3d showed significant survival differences (p=1.05 *×* 10^−4^), indicating that our model effectively distinguished high-risk and low-risk patients: genes such as hsa-miR-148a-3p, AKR1E2, and PCDHB2 were upregulated in the high-risk groups (Q1–Q2), whereas hsa-miR-363-3p, DUSP15, and GSTM4 were upregulated in the low-risk groups (Q3–Q4), consistent with the directions revealed by the above analysis. When patients were instead stratified by the median predicted risk (Figure 3e), the high-risk group again exhibited significantly worse survival (p=1.42 *×* 10^−3^), with AKR1E2, ARMC1, SDHAF4, TMEM242, and PCDHB2 upregulated and NDUFV2-AS1, GSTM4, DUSP15, SUSD3, TPRG1, and RFX2 downregulated in the high-risk group, further confirming that the risk signals captured by the model were aligned with actual molecular changes. Consistently, the distribution of predicted risk scores across breast cancer molecular subtypes (Figure 3h) showed that Basal and Her2 subtypes had the highest risk, LumA and LumB subtypes had markedly lower risk, and LumA had the lowest risk, matching known biological characteristics of breast cancer subtypes [53].

To validate the effectiveness of these 14 significant genes, we built a multivariable CoxPH model on the training set using their expression levels and evaluated its performance on the test set. Figure 3g illustrated that CoxPH significantly separated high-risk and low risk patients (HR=3.60, p=0.0149, C-Index=0.6998), indicating good generalization of the survival-related genes identified by LaCONIC. To further explore potential regulatory relationships, we constructed a gene interaction network based on a known ceRNA network (Figure 3f), in which hsa-miR-148a-3p, hsa-miR-363-3p, hsa-miR-365a-3p, TMEM242 and GSTM4 formed a core interaction cluster connected to risk genes such as ARMC1 and TMEM242. This finding suggested synergistic effects in tumor progression and prognosis. Consistently, multi-gene survival analyses in Figure 3i showed that several gene combinations (e.g., hsa-miR-148a-3p + hsa-miR-363-3p + GSTM4 + TMEM242, TMEM242 + ARMC1 + hsa-miR-365a-3p) significantly stratified patients into high-risk and low-risk groups (p¡0.05), with markedly poorer survival in the high-expression groups, demonstrating that the core genes identified by LaCONIC had independent prognostic value and exhibited synergistic risk effects, thereby supporting the biological plausibility and clinical relevance of our model’s predictions.

#### 2) Enrichment analysis

To investigate the biological functions of these 14 genes identified by LaCONIC, we performed pathway enrichment analysis across Gene Ontology Biological Process (GOBP), KEGG, WikiPathways and Reactome. At the GOBP level (Figure 4a), these genes were primarily enriched in dephosphorylation, glial cell differentiation, mitochondrion organization, synapse assembly, and DNA replication, implicating tumor metabolism, neuroendocrine-like transdifferentiation, and cell-cycle activity, with TMEM242, ARMC1, PCDHB2, and SDHAF4 recurrent across multiple terms [54], [55]. KEGG analysis (Figure 4b) indicated that GSTM4, hsa-miR-365a-3p, hsa-miR-363-3p, and hsa-miR-148a-3p were mainly involved in chemical carcinogenesis, AMPK signaling, and AGE–RAGE signaling, suggesting that LaCONIC-identified genes may affect survival via energy metabolism and signal transduction [56]. WikiPathways enrichment (Figure 4c) further highlighted miRNA-target regulation pathways, BDNF signaling, and the ‘CDK4/6 inhibitors in breast cancer’ pathway, indicating potential roles in tumor signaling, proliferation, and cell-cycle control via the CDK4/6–Cyclin D–Rb axis [57]. Reactome analysis (Figure 4d) supported these findings by revealing enrichment in NMDA receptor signaling, adherens junctions, Wnt signaling, and apoptosis, with genes such as RFX2 and GSTM4 shared across multiple modules [58]. Together, these results showed that LaCONIC-identified genes were tightly linked to tumor-related biological processes, energy metabolism, and neural-associated regulatory networks, reinforcing the biological plausibility of the model’s risk predictions.

**Fig. 4.**
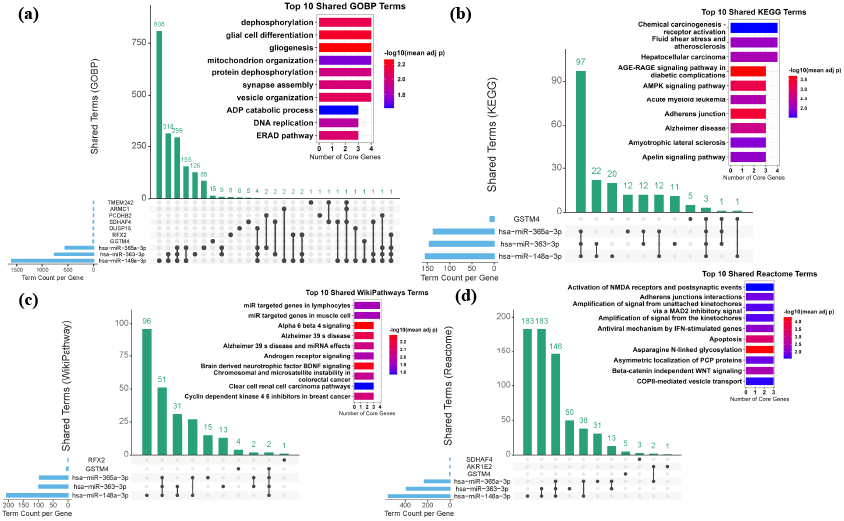
Enrichment analysis of genes significantly related to survival.

### E. Latent space visualization analysis

To further characterize the latent structure learned by LaCONIC for survival risk and molecular phenotypes, we performed UMAP-based visualization and correlation analysis on the BRCA test set. As shown in Figure 5(a), the survival-related latent vector *Z*_surv_ clearly separated the four risk quartiles (Q1–Q4): high-risk patients (Q1, Q2) clustered at one end of the latent space, forming a ‘poor prognosis’ region, whereas low-risk patients (Q3, Q4) were more dispersed and formed several subclusters, consistent with a continuous risk gradient and an underlying risk evolution trajectory. The middle and right panels showed that both *Z*_surv_ and *Z*_cls_ separated breast cancer subtypes, with sharper clusters for *Z*_cls_. Jointly examining the left and middle panels further revealed that LumA samples concentrated in low-risk regions, whereas Basal samples were enriched in high-risk regions, indicating that LaCONIC aligned survival risk with subtype information and encoded both continuous risk gradients and subtype-specific molecular patterns in its latent space.

**Fig. 5.**
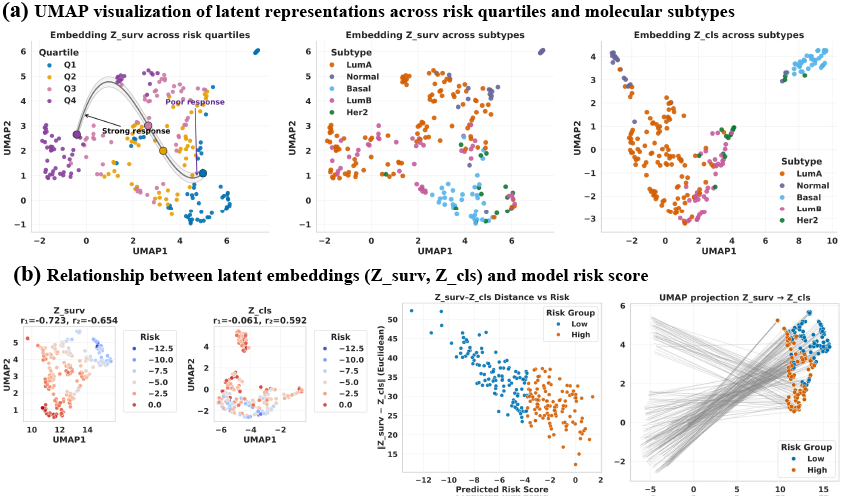
Integrated survival and subtype related latent representations reveal risk-associated structure in the embedding space.

Figure 5(b) summarized the relationship between the survival representation *Z*_surv_, the classification representation *Z*_cls_, and the predicted risk scores. The left panels showed a strong negative correlation between *Z*_surv_ and risk (r = −0.723), indicating that latent position encoded a clear survival risk gradient, whereas *Z*_cls_ showed only weak correlation (r = −0.061). The middle panel showed that the Euclidean distance ∥ *Z*_surv_ − *Z*_cls_ ∥ decreased with increasing risk, implying more compact and consistent latent representations for high-risk patients. The right UMAP projection further indicated that low-risk samples occupied a broader region between *Z*_surv_ and *Z*_cls_, while high-risk samples converged, suggesting a tighter fusion of survival and subtype information. Together, these results indicated that LaCONIC effectively integrated survival signals and molecular features at the representation level.

### F. Ablation study

To comprehensively evaluate the effectiveness of the LaCONIC model, we conducted ablation experiments on all cancer datasets from the following three aspects, and evaluated <u>the performance using Harrell’s C-Index.</u>

#### 1) Architectural components

To assess the contribution of each architectural component in LaCONIC, we conducted ablation experiments by removing heterogeneous graph self-supervised pretraining (w/o HGT), ACRL (w/o ACRL), GCIL (w/o GCIL), and gene graph features (w/o graph feature). In the w/o HGT setting, pretrained graph embeddings were replaced by random embeddings to retain gene-level features in both modal; in the w/o graph feature setting, only gene expression features were used. As shown in Figure 6(a), the whole LaCONIC achieved the best performance across all datasets, with particularly large gains on CESC and SARC. Removing HGT or graph features caused substantial performance drops, especially on BRCA and UCEC, indicating the critical role of structured graph representations and the detrimental effect of random graph features. Eliminating ACRL or GCIL also degraded performance on most datasets, suggesting that these modules effectively capture multi-omics representations and cross-omics dependencies. Overall, LaCONIC’s performance depends on the synergy of these modules.

**Fig. 6.**
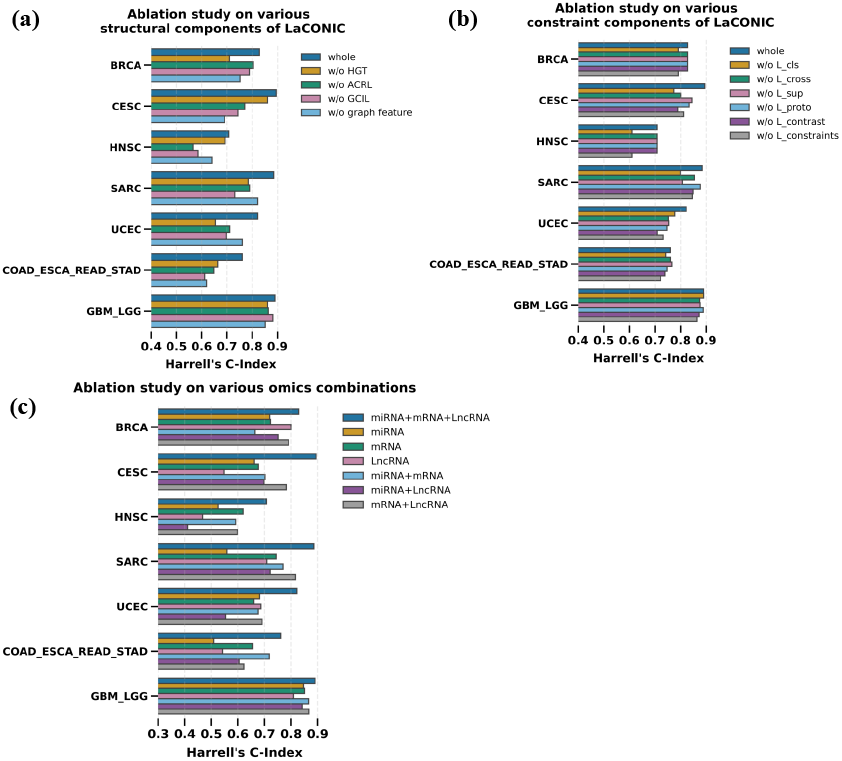
Ablation experiment results based on Harrell’s C-Index from different perspectives.

#### 2) Label-aware constraint

To assess the effect of the label-aware constraint mechanism, we ablated subtype classification loss (w/o *L*_cls_), cross-task alignment loss (w/o *L*_cross_), supervised contrastive loss (w/o *L*_sup_), prototype contrastive loss (w/o *L*_proto_), the combined contrastive term (*L*_contrast_ = *L*_cross_+*L*_sup_+*L*_proto_), and the full constraint block (*L*_constraints_ = *L*_cls_+*L*_contrast_). As shown in Figure 6(b), the complete label-aware constraint consistently yielded the best performance, and removing *L*_constraints_ caused large drops, especially on COAD ESCA READ STAD and GBM LGG. *L*_cls_ was critical on most datasets and was the only non-survival loss with a strong impact on BRCA and HNSC. The impact of individual contrastive losses was cancer-specific, while removing the combined *L*_contrast_ produced larger performance degradations than ablating any single contrastive term, underscoring the importance of multi-level contrastive constraints for LaCONIC’s generalization and survival prediction accuracy.

#### 3) Multi-omics combinations

To quantify the contribution of different omics, we performed multi-omics ablations using miRNA, mRNA, and lncRNA individually, in all pairwise combinations, and in the full three-omics setting. As shown in Figure 6(c), integrating miRNA+mRNA+lncRNA consistently improved performance across cancers, with marked gains on CESC and SARC, while pairwise combinations generally outperformed single-omics inputs. Among single modalities, mRNA showed the most robust performance, miRNA was particularly informative in nervous system tumors (e.g., GBM LGG), and lncRNA added value in cancers such as BRCA. To avoid hyperparameter bias, we further retuned LaCONIC on the CESC dataset for each omics combination; as reported in Supplementary Table S3, the miRNA+mRNA+lncRNA setting achieved the best results on all metrics (Harrell’s C-Index = 0.8943, iAUC = 0.9093, IPCW C-Index = 0.8980), with mRNA alone ranking second and miRNA alone yielding the highest IPCW C-Index. Although lncRNA alone performed worst, its combination with mRNA produced clear gains. Overall, these results indicated complementary and synergistic effects among miRNA, mRNA, and lncRNA and showed that multi-omics integration substantially improved the accuracy and robustness of survival prediction.

## V. Conclusion

Accurate cancer survival prediction is essential for guiding treatment and reducing mortality. To overcome limitations in multi-omics integration and network modeling, we proposed LaCONIC integrating heterogeneous graph self-supervised pretraining, adaptive cross-omics representation learning, graph-guided cross-omics interaction modules, and multi-level label-aware constraints to collaboratively capture molecular network topology and subtype-discriminative features. Experiments on multiple cancer datasets showed that LaCONIC consistently outperforms existing survival models, demonstrating strong representational and generalization capabilities. Interpretability analysis on breast cancer highlighted survival-associated genes that were enriched in pathways related to signal transduction, energy metabolism, cell-cycle regulation, and the “CDK4/6 inhibitors in breast cancer”, supporting their mechanistic relevance. Latent space visualizations further revealed a smooth risk gradient and clearly separated molecular subtypes, underscoring the interpretability and biological consistency of the learned representations.

While LaCONIC performs well in survival prediction and interpretability, it has several limitations. First, model training relies on high-quality multi-omics paired data, so missing modalities and batch effects may hinder generalization. Second, although interpretability analyses provide some feature interpretation, experimental validation of the identified regulators and pathways is still required. In future work, we plan to explore cross-cancer transfer learning and dynamic graph modeling to capture temporal biological processes such as tumor evolution and treatment response, and to integrate single-cell omics and spatial transcriptomics data to more comprehensively characterize cellular heterogeneity and the tumor microenvironment.

## Acknowledgments

This work is supported by the National Natural Science Foundation of China (#62372165 and #62032007).

