## Supplementary Materials for "LaCONIC: A Label-Aware and Graph-Guided Contrastive Multi-Omics Collaborative Learning Model for Cancer Risk Prediction"

November 26, 2025

### 1 Supplementary Material Section 1: Dataset and preprocessing

For the expression data of all cancers, we implemented a strict quality control and feature screening process: (1) Eliminating genes that were missing in more than 10% of the samples; (2) Removing samples with gene deletions exceeding 50%; (3) Retaining genes with an expression level greater than 1 in at least 20% of the samples; (4) Performing log2 transformation and standardization on the retained expression matrix; (5) Adopting univariate Cox regression analysis to screen genes significantly associated with Overall Survival (OS), so as to retain the molecular characteristics that were most statistically significant for prognostic events.

Table S1: The number of edges/nodes and their sources of various relational data

| | Number of edges/nodes | Source |
|---|---|---|
| miRNA | 701 | miRBase v22 Kozomara et al. (2019) |
| mRNA | 17,236 | R package `biomaRt` [HGNC Symbol] Durinck et al. (2005) |
| lncRNA | 5,423 | R package `biomaRt` [HGNC Symbol] |
| disease | 867 | MeSH subject terms Lipscomb (2000) |
| miRNA-mRNA | 1,288,514 | miRtarbase v10.0 Cui et al. (2025) ∪ Tarbase v9.0 Skoufos et al. (2024) |
| miRNA-disease | 28,755 | HMDD v4.0 Cui et al. (2024) |
| miRNA-lncRNA | 5,495 | starbase Li et al. (2014) |
| mRNA-mRNA | 12,335,080 | STRING PPI v12.0 Szklarczyk et al. (2023) |
| mRNA-disease | 8,506,622 | DisGeNET Piñero et al. (2020) |
| lncRNA-disease | 5,443 | LncRNADisease v3.0 ∪ v2.0 Bao et al. (2019) |
| miRNA-miRNA | 83,086 | Zhu et al Zhu et al. (2013) |

Table S2: An overview of cancer-specific relational and ceRNA network data

| | BRCA | CESC | HNSC | SARC | UCEC | COAD_ESCA _READ_STAD | GBM_LGG |
|---|---|---|---|---|---|---|---|
| miRNA | 116 | 78 | 118 | 174 | 70 | 69 | 193 |
| mRNA | 4129 | 1849 | 1537 | 2400 | 2931 | 1554 | 6285 |
| lncRNA | 141 | 120 | 55 | 90 | 165 | 43 | 376 |
| disease | 866 | 845 | 857 | 860 | 853 | 850 | 867 |
| miRNA-mRNA | 90,638 | 15,328 | 20,068 | 81,984 | 34,131 | 14,470 | 235,423 |
| miRNA-disease | 8062 | 4909 | 6361 | 7850 | 3754 | 4169 | 11,880 |
| miRNA-lncRNA | 165 | 27 | 65 | 141 | 117 | 44 | 570 |
| mRNA-mRNA | 472,621 | 111,551 | 73,206 | 197,335 | 226,467 | 77,518 | 1,047,725 |
| mRNA-disease | 2,176,118 | 972,872 | 799,323 | 1,285,422 | 1,526,232 | 836,933 | 3,307,503 |
| lncRNA-disease | 605 | 354 | 167 | 507 | 584 | 321 | 1,165 |
| miRNA-miRNA | 4336 | 1450 | 2600 | 9204 | 1311 | 1005 | 11,149 |
| ceRNA | 561,289 | 127,028 | 95,099 | 286,299 | 259,070 | 91,982 | 1,280,772 |

#### 2 Supplementary Material Section 2: Topological Feature Initialization

The initial node-level topological features is constructed from the betweenness centrality of the ceRNA network. Betweenness centrality is defined as

$$C_B(v) = \frac{1}{(G-1)(G-2)} \sum_{\substack{s,t \in \mathcal{V} \\ s \neq t \neq v}} \frac{\sigma_{st}(v)}{\sigma_{st}}, \tag{1}$$

where $\sigma_{st}$ denotes the number of shortest paths between nodes $s$ and $t$, and $\sigma_{st}(v)$ is the number of those paths that pass through node $v$. This metric reflects the 'bridge' role of a node in signal transduction and regulatory pathways, and thus serves as an important topological feature for identifying key regulatory genes and hub miRNAs.

Given the high computational cost of exact betweenness computation in large-scale biological networks, we adopt an adaptive sampling approximation based on Brandes' algorithm Brandes (2001). Specifically, the number of sampled source nodes $k$ is determined dynamically according to the number of nodes $G$ and network density $\rho$:

$$k = \begin{cases} \min\big(50 \log_{10}(G+1) \times 2,\, k_{\max}\big), & \rho < 0.01, \\ \max\big(50 \log_{10}(G+1)/2,\, k_{\min}\big), & \rho \geq 0.01. \end{cases} \tag{2}$$

We then randomly sample $k$ source nodes from the graph and use breadth-first search to compute node dependencies, which are accumulated and averaged to obtain the approximate betweenness centrality:

$$\tilde{C}_B(v) = \frac{1}{k(G-1)(G-2)} \sum_{s=1}^{k} \delta_v^{(s)}, \tag{3}$$

where $\delta_v^{(s)}$ denotes the dependency of node $v$ with respect to the $s$-th sampled source node. Finally, the resulting values are assembled into the topological feature matrix $X_{\text{topo}} = \tilde{C}_B(v) \in \mathbb{R}^{g \times 1}$, which is fed into the GCIL module to enhance the network awareness of node representations.

### 3 Supplementary Material Section 3: Experimental setup

#### 3.1 Metrics

To systematically evaluate the model's performance in cancer survival prediction and risk stratification tasks, this study used various clinical survival analysis and statistical testing metrics, including Harrell's C-Index, iAUC, IPCW C-Index, and risk stratification, to comprehensively characterize the model's predictive ability and stability across different levels.

**1) Harrell's Concordance Index (Harrell's C-Index)** Harrell et al. (1982):
The Harrell's C-Index is used to assess the consistency of the model's ranking of individual survival times Harrell et al. (1982), defined as follows:

$$C_{\text{Harrell}} = \frac{1}{N_{\text{comp}}} \sum_{i,j} \mathbb{I}[(t_i < t_j) \wedge (r_i > r_j)] \,, \tag{4}$$

where $t_i$ is the survival time of sample $i$; $r_i$ is the model-predicted risk score; and $N_{\text{comp}}$ is the number of comparable sample pairs, which only include pairs of samples with an event (death), excluding censored samples.

**2) Integrated AUC (iAUC)** Heagerty et al. (2000):
The Integrated AUC (iAUC) is the integral of the time-dependent ROC curve AUC, used to evaluate the model's discriminatory ability at different follow-up time points Heagerty et al. (2000), defined as:

$$\text{iAUC} = \frac{1}{T} \int_0^T \text{AUC}(t)\, dt, \tag{5}$$

where AUC($t$) represents the instantaneous predictive performance at time $t$, calculated based on cumulative dynamic AUC. The time evaluation points are selected as the 25%, 50%, and 75% percentiles of the survival time distribution.

**3) Inverse Probability of Censoring Weighting C-Index (IPCW C-Index)** Uno et al. (2011):
The IPCW C-Index is used to reduce the bias caused by censored data Uno et al. (2011), defined as:

$$C_{\text{IPCW}} = \sum_{i<j} w_{ij}\, \mathbb{I}[(t_i < t_j) \wedge (\hat{r}_i > \hat{r}_j)] \,, \tag{6}$$

where $w_{ij}$ is the censoring adjustment weight based on individual survival probability estimates.

**4) Kaplan–Meier Risk Stratification Analysis** Pencina et al. (2008):
Kaplan–Meier risk stratification divides samples into high-risk and low-risk groups based on the model's

predicted median risk, and Kaplan–Meier (KM) survival curves Pencina et al. (2008); Kaplan and Meier (1958) are plotted to show survival differences between the groups. The significance of group differences is analyzed using the Log-rank test Mantel et al. (1966), and the p-value is used to assess statistical differences between the survival curves of high- and low-risk groups. To quantify the mortality risk of the high-risk group relative to the low-risk group, the Hazard Ratio (HR) and its 95% confidence interval are calculated using the Cox proportional hazards model. HR and its confidence interval are derived when both groups experience events, reflecting the model's discriminatory ability and stability in survival risk stratification.

#### 3.2 Dataset division and sampling

To ensure the representativeness and stability of multi-modal cancer data during the training and validation phases, this study adopted a strategy combining stratified partitioning and balanced batch sampling to mitigate issues arising from event censoring and imbalanced subtype distribution in survival analysis. Data division was based on the combination of survival event status and subtype labels, ensuring a balanced distribution of categories and event ratios across subsets. The original data was first split into an 80% training set and a 20% test set, with 20% of the training set further reserved as a validation set. The final data partitioning ratio was 64% for training, 16% for validation, and 20% for testing.

During the training phase, to address the issue of event sample scarcity, we designed a balanced batch sampler to ensure that each batch contains at least $k$ event samples, stabilizing gradient estimates. Let the batch size be $B$, the number of event samples be $N_1$, and the total number of training samples be $N$. The minimum number of event samples was calculated as $k = \left\lceil \frac{N_1}{N/B} \right\rceil$. This mechanism ensured an adequate proportion of event samples in each batch, enhancing the stability of Cox loss and time-dependent AUC. With this stratified partitioning and balanced batch strategy, the model's robustness in sample distribution and risk estimation was significantly improved.

#### 3.3 Training strategy of LaCONIC

To achieve stable convergence and task collaboration in the multi-task multi-omics model, LaCONIC employed a strategy combining stage-wise weight ramp-up and smooth learning rate scheduling during training. Specifically, the training was divided into three stages: In Stage 0, the model optimized only survival prediction and classification tasks to stabilize the foundational feature space; in Stage 1, the cross-task loss $\mathcal{L}$cross and supervised contrastive loss $\mathcal{L}$sup were gradually introduced, with their weights increasing linearly to align risk and subtype features collaboratively; in Stage 2, the prototype contrastive constraint $\mathcal{L}_{\text{proto}}$ was enabled, further enhancing the boundary clarity between different subtypes. This progressive scheduling mechanism effectively prevented gradient conflicts during the early stages of multi-task training, enabling the model to learn high-level feature associations while converging stably.

In addition, the AdamW optimizer was used with learning rate warmup and cosine annealing strategies to ensure smooth parameter updates and stable learning rate changes. The model's performance was evaluated after each iteration using the validation set's Harrell's C-Index and a composite loss function, with the best weights saved when performance improves. Early stopping was triggered if no improvement was observed over several consecutive iterations (default is 50) to prevent overfitting. This training strategy enhanced both task collaboration and generalization, providing an efficient and reliable optimization paradigm for cancer multi-omics risk prediction.

#### 3.4 Implementation details

To evaluate the performance of LaCONIC, we compared it against 14 existing methods, including three classical survival models (Coxnet, CoxBoost Weyer and Binder (2015), RSF), four deep discrete-time models (DeepHit, Transformer Survival, Trans-STG, SurvTRACE), and seven deep continuous-time models (DeepSurv, Cox-nnet, DCAP, HFBSurv, CAMR, FGCNSurv, PCLSurv). Three classical methods were implemented using the Python library `sksurv` Pölsterl (2020), DeepSurv and DeepHit were implemented with `PyCox` Kvamme et al. (2019), and the remaining deep models were implemented in `PyTorch`. Transformer Survival, Trans-STG, and DCAP were reimplemented according to their original papers, whereas SurvTRACE, HFBSurv, CAMR, FGCNSurv, and PCLSurv were run using the official GitHub implementations released by the authors. To ensure a fair comparison, all methods except FGCNSurv and PCLSurv (which were inherently designed to use only miRNA and mRNA) were trained on the same multi-omics inputs. Moreover, except for HFBSurv, CAMR, FGCNSurv, and PCLSurv (which employed a specific encoder for each omics), the remaining baselines took the concatenated three-omics gene expression matrix as input.

Table S3: Comparison of survival prediction performance for different omics combinations on the CESC dataset

| Omics combination | Harrell's C-Index | iAUC | IPCW C-Index |
|---|---|---|---|
| miRNA+mRNA+lncRNA | **0.8943** | **0.9093** | **0.8980** |
| mRNA | <u>0.8792</u> | <u>0.9064</u> | 0.7790 |
| mRNA+lncRNA | 0.8491 | 0.8611 | 0.8407 |
| miRNA+mRNA | 0.8302 | 0.8621 | 0.7600 |
| miRNA | 0.8151 | 0.8193 | <u>0.8565</u> |
| miRNA+lncRNA | 0.7849 | 0.7761 | 0.8237 |
| lncRNA | 0.7132 | 0.7211 | 0.6481 |

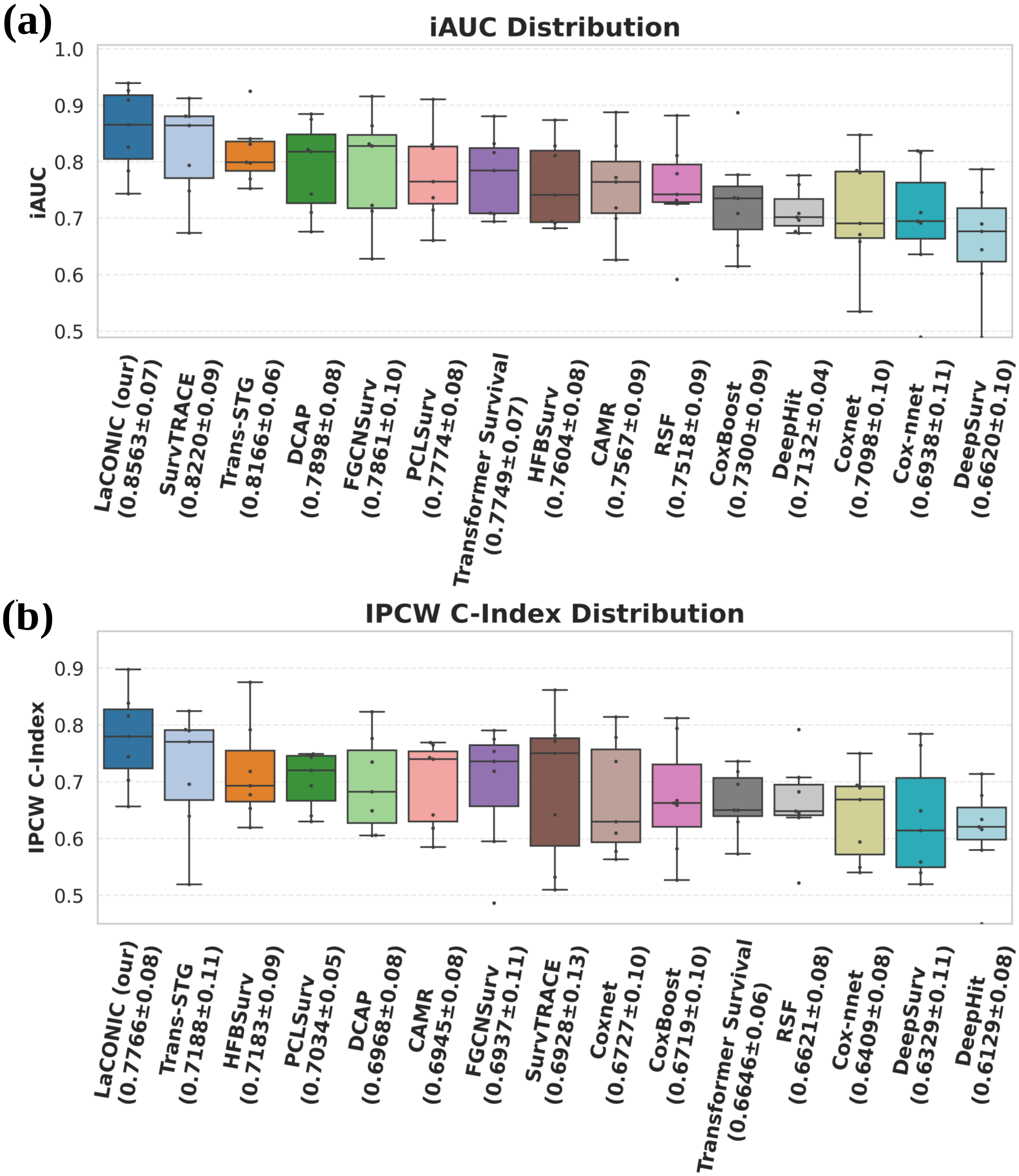

Figure S1: Comparison of the distributions of iAUC and IPCW C-index for all models on seven cancer datasets

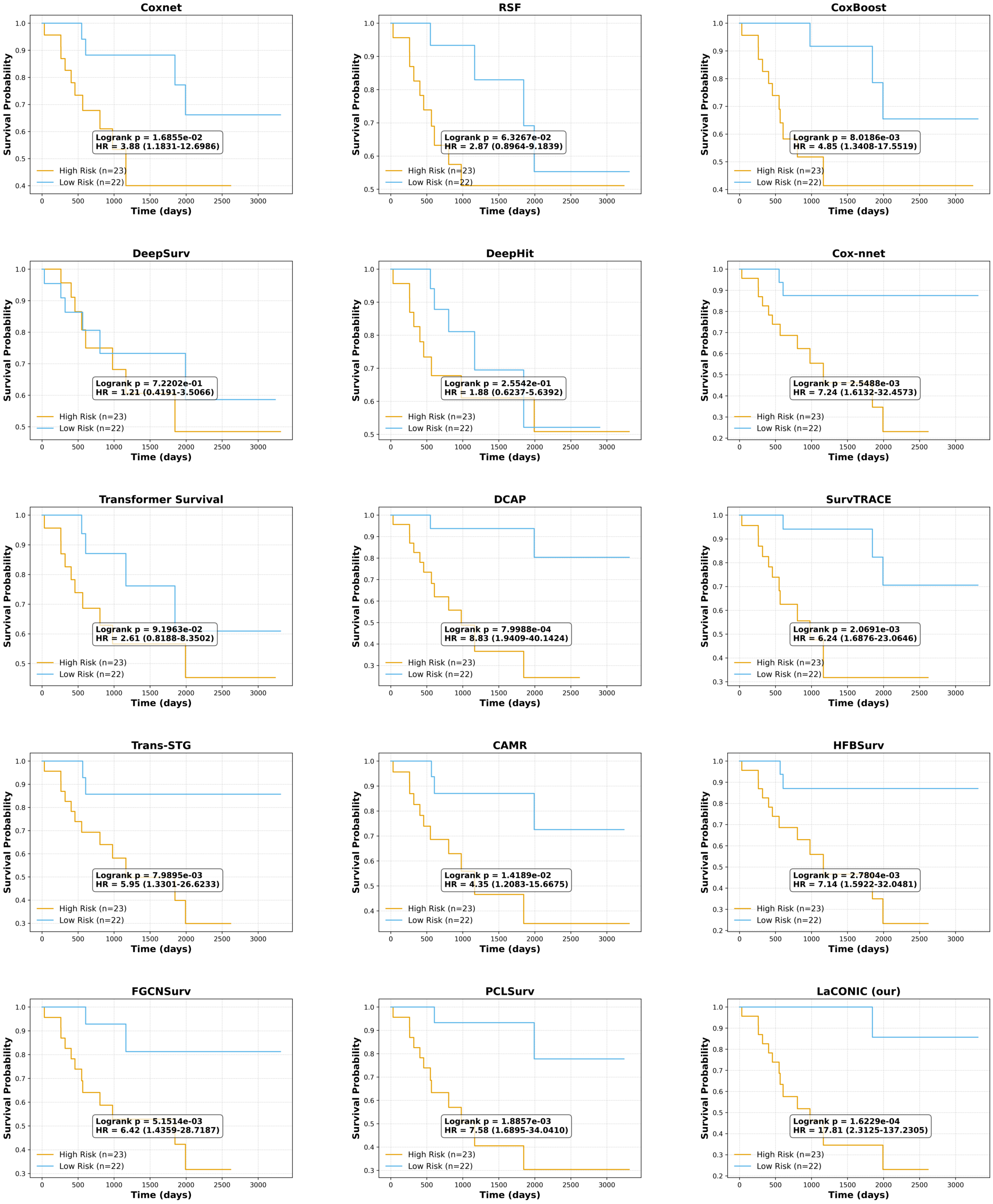

Figure S2: The Kaplan-Meier risk stratification curves of all models on the SARC dataset

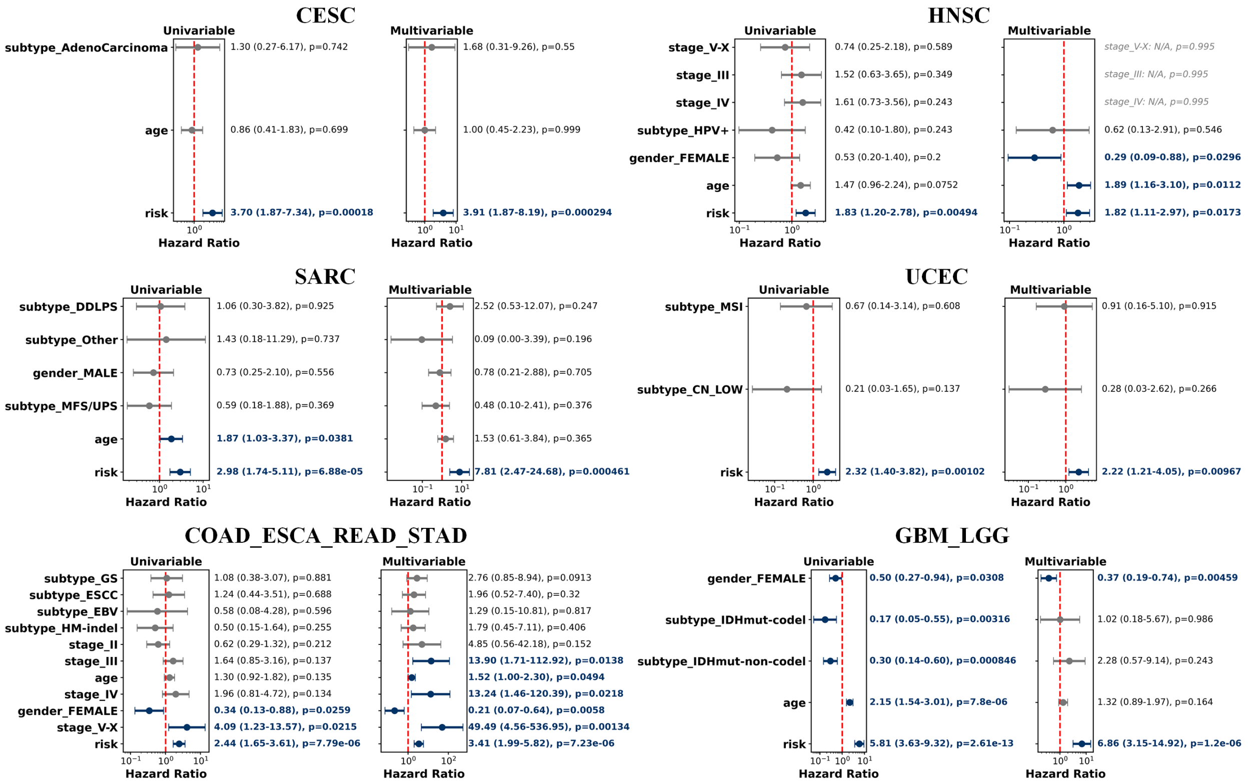

Figure S3: Univariable and multivariable Cox proportional hazards analysis of LaCONIC-predicted risk and clinical covariates across four single-cancer and two pan-cancer datasets

#### 3.5 Hyperparameter settings

To ensure a fair and efficient hyperparameter search, this study employed Optuna to tune the hyperparameters of LaCONIC and all baseline models, with the random seed uniformly fixed at 42 to facilitate reproducibility. Considering that the dimensionality and search space of hyperparameters differ across models, we assigned an appropriate number of trials to each model (Detailed setting in Table S4).

Table S4: Number of Optuna hyperparameter search trials for each method

| Model | trial number |
|---|---|
| LaCONIC | 200 |
| Coxnet | 50 |
| RSF | 150 |
| CoxBoost | 200 |
| DeepSurv | 200 |
| DeepHit | 200 |
| Cox-nnet | 150 |
| Transformer Survival | 200 |
| DCAP | 100 |
| SurvTRACE | 200 |
| TransSTG | 200 |
| CAMR | 75 |
| HFBSurv | 75 |
| FGCNSurv | 75 |
| PCLSurv | 20 |

The hyperparameters of LaCONIC hyperparameters includes batch size, attention/classifier/survival dropout rates, EMA momentum, training epochs, hidden layer dimensions and depth of classifier/survival predictor, intermediate dimension in GCIL module, learning rate, number of attention heads and output dimensions in ACRL module, temperatures for cross/prototype/supervised contrastive losses, and the corresponding loss weights (cross, prototype, supervised, and classfication). Under the miRNA+mRNA+lncRNA multi-omics combination, the optimal hyperparameter settings of LaCONIC across all cancer types were summarized in Table S5, whereas the optimal hyperparameters under different omics combinations for the CESC dataset were presented in Table S6. The optimal hyperparameter settings of the other comparison methods were summarized in Tables S7–Table S20.

Table S5: Best hyperparameters of LaCONIC across cancers

| param | BRCA | CESC | HNSC | SARC | UCEC | COAD _ESCA _READ _STAD | GBM _LGG |
|---|---|---|---|---|---|---|---|
| best_c_index | 0.828248031 | 0.894339623 | 0.707492795 | 0.884422111 | 0.821782178 | 0.760298661 | 0.889316988 |
| batch_size | 16 | 16 | 24 | 16 | 24 | 24 | 8 |
| dropout_att1 | 0.068555596 | 0.068563974 | 0.080922912 | 0.044009127 | 0.035735726 | 0.031921386 | 0.004273933 |
| dropout_att2 | 0.027318465 | 0.045832206 | 0.028476266 | 0.027819807 | 0.008836704 | 0.020991902 | 0.001986249 |
| dropout_cls | 0.039415717 | 0.276581389 | 0.218186753 | 0.19035103 | 0.183115986 | 0.387294427 | 0.19984428 |
| dropout_surv | 0.135118334 | 0.033940272 | 0.041874682 | 0.146065789 | 0.060142412 | 0.280037287 | 0.080410038 |
| ema_m | 0.929585169 | 0.967138415 | 0.980374947 | 0.946690413 | 0.901331893 | 0.88883725 | 0.96526094 |
| epochs | 200 | 200 | 200 | 200 | 200 | 200 | 200 |
| hidden_dims | [256, 224] | [128, 96] | [128] | [128] | [224, 160] | [160, 160] | [64, 64] |
| hidden_n_layers | 2 | 2 | 1 | 1 | 2 | 2 | 2 |
| inter_model | 16 | 64 | 32 | 16 | 64 | 32 | 16 |
| lr | 0.001159917 | 0.002241198 | 0.003971425 | 0.000998748 | 0.000239019 | 0.000740467 | 0.000216812 |
| num_heads | 2 | 1 | 2 | 1 | 2 | 2 | 1 |
| o_dims | [16, 16, 16] | [32, 32, 32] | [16, 16, 16] | [16, 16, 16] | [16, 16, 16] | [16, 16, 16] | [32, 32, 32] |
| temp_cross | 0.126960286 | 0.109216443 | 0.084026541 | 0.074625476 | 0.073747873 | 0.137328299 | 0.101770808 |
| temp_proto | 0.113330171 | 0.071744504 | 0.062913029 | 0.105313326 | 0.102274767 | 0.081560676 | 0.066410544 |
| temp_sup | 0.195456452 | 0.19755224 | 0.152274852 | 0.194525396 | 0.12830436 | 0.165778486 | 0.206739436 |
| weight_cross | 0.160332632 | 0.109301884 | 0.055345172 | 0.086849955 | 0.178947764 | 0.179448773 | 0.170426181 |
| weight_decay | 5.95E-05 | 6.19E-05 | 1.47E-07 | 3.21E-07 | 1.78E-07 | 8.70E-07 | 1.27E-05 |
| weight_proto | 0.006468953 | 0.091604206 | 0.087769499 | 0.017355726 | 0.094696382 | 0.117411948 | 0.071058932 |
| weight_sup | 0.182471357 | 0.240996404 | 0.19505594 | 0.232218319 | 0.187838378 | 0.204075648 | 0.181349014 |
| weight_task | 0.5 | 0.7 | 0.7 | 0.7 | 0.1 | 0.5 | 0.1 |

Table S6: Best hyperparameters of LaCONIC under various omic combinations on CESC data

| Hyperparameters | miRNA | mRNA | lncRNA | miRNA+mRNA | miRNA+lncRNA | mRNA+lncRNA | miRNA+mRNA+lncRNA |
|---|---|---|---|---|---|---|---|
| best_c_index | 0.81509434 | 0.879245283 | 0.713207547 | 0.830188679 | 0.78490566 | 0.849056604 | 0.894339623 |
| batch_size | 16 | 16 | 16 | 16 | 16 | 16 | 16 |
| dropout_att1 | 0.037397771 | 0.093294115 | 0.023296206 | 0.017024219 | 0.060256102 | 0.028225835 | 0.068563974 |
| dropout_att2 | 0.064042786 | 0.091092466 | 0.043392725 | 0.000869357 | 0.02952621 | 0.008097519 | 0.045832206 |
| dropout_cls | 0.245826332 | 0.16202801 | 0.270829515 | 0.04161505 | 0.08635746 | 0.016094939 | 0.276581389 |
| dropout_surv | 0.027155327 | 0.086184121 | 0.195424328 | 0.204044094 | 0.06522064 | 0.254364128 | 0.033940272 |
| ema_m | 0.91025237 | 0.989904877 | 0.981549092 | 0.922221804 | 0.978033327 | 0.924601457 | 0.967138415 |
| epochs | 200 | 200 | 200 | 200 | 200 | 200 | 200 |
| hidden_dims | [160, 96] | [160] | [192] | [224] | [160, 128] | [224] | [128, 96] |
| hidden_n_layers | 2 | 1 | 1 | 1 | 2 | 1 | 2 |
| inter_model | 64 | 64 | 32 | 32 | 16 | 16 | 64 |
| lr | 0.001941776 | 0.003172307 | 0.00355262 | 0.001000495 | 0.004636073 | 0.000655382 | 0.002241198 |
| num_heads | 2 | 2 | 1 | 1 | 2 | 1 | 1 |
| o_dims | [32, 32, 32] | [32, 32, 32] | [16, 16, 16] | [16, 16, 16] | [32, 32, 32] | [16, 16, 16] | [32, 32, 32] |
| temp_cross | 0.122987302 | 0.075557684 | 0.092093164 | 0.092477109 | 0.071703794 | 0.102675691 | 0.109216443 |
| temp_proto | 0.103485087 | 0.090577273 | 0.101900742 | 0.100800204 | 0.113718338 | 0.118849466 | 0.071744504 |
| temp_sup | 0.217726647 | 0.200140365 | 0.125938892 | 0.155536319 | 0.155839416 | 0.166732911 | 0.19755224 |
| weight_cross | 0.170408792 | 0.100058607 | 0.054380153 | 0.129261241 | 0.136925993 | 0.051473682 | 0.109301884 |
| weight_decay | 1.25E-05 | 3.16E-07 | 2.14E-05 | 3.56E-06 | 5.24E-06 | 2.24E-07 | 6.19E-05 |
| weight_proto | 0.00235547 | 0.059342691 | 0.033624805 | 0.005207007 | 0.056940275 | 0.054306061 | 0.091604206 |
| weight_sup | 0.279779056 | 0.302003685 | 0.329361449 | 0.155685673 | 0.303497662 | 0.293145995 | 0.240996404 |
| weight_task | 0.9 | 0.3 | 0.3 | 1 | 1 | 0.3 | 0.7 |

Table S7: Best hyperparameters of CAMR across cancers

| Hyperparameters | BRCA | CESC | HNSC | SARC | UCEC | COAD _ESCA _READ _STAD | GBM _LGG |
|---|---|---|---|---|---|---|---|
| best_score | 8.041339e-01 | 0.764151 | 6.001441e-01 | 8.178392e-01 | 7.186469e-01 | 0.701081 | 0.842032 |
| lr | 1.809691e-04 | 0.000834 | 2.170219e-03 | 3.083498e-03 | 4.169858e-03 | 0.000122 | 0.000259 |
| lambda_reg | 6.783867e-05 | 0.000001 | 1.525521e-05 | 1.484840e-04 | 5.359932e-07 | 0.000008 | 0.000001 |
| weight_decay | 3.187451e-07 | 0.000006 | 1.817557e-07 | 2.065507e-07 | 1.042297e-07 | 0.000012 | 0.000011 |

Table S8: Best hyperparameters of FGCNSurv across cancers

| Hyperparameters | BRCA | CESC | HNSC | SARC | UCEC | COAD _ESCA _READ _STAD | GBM _LGG |
|---|---|---|---|---|---|---|---|
| best_score | 0.808961 | 0.826415 | 6.037464e-01 | 8.161209e-01 | 0.689256 | 6.975484e-01 | 0.865520 |
| k | 8.000000 | 8.000000 | 7.000000e+00 | 4.000000e+00 | 10.000000 | 1.000000e+01 | 4.000000 |
| lr | 0.000306 | 0.000433 | 4.770725e-04 | 9.717156e-04 | 0.000448 | 1.540745e-03 | 0.000842 |
| weight_decay | 0.000027 | 0.000071 | 1.456504e-07 | 3.167275e-07 | 0.000004 | 1.111996e-07 | 0.000004 |

Table S9: Best hyperparameters of HFBSurv across cancers

| Hyperparameters | BRCA | CESC | HNSC | SARC | UCEC | COAD _ESCA _READ _STAD | GBM _LGG |
|---|---|---|---|---|---|---|---|
| best_score | 7.554134e-01 | 0.807547 | 0.647695 | 8.115578e-01 | 0.686469 | 0.669156 | 0.832574 |
| lr | 7.475993e-04 | 0.000180 | 0.003675 | 2.696155e-04 | 0.000543 | 0.000245 | 0.000101 |
| lambda_reg | 7.669581e-07 | 0.000101 | 0.000028 | 4.911117e-05 | 0.000001 | 0.000177 | 0.000083 |
| weight_decay | 5.987475e-06 | 0.000049 | 0.000002 | 7.921335e-07 | 0.000014 | 0.000027 | 0.000002 |

Table S10: Best hyperparameters of PCLSurv across cancers

| Hyperparameters | BRCA | CESC | HNSC | SARC | UCEC | COAD _ESCA _READ _STAD | GBM _LGG |
|---|---|---|---|---|---|---|---|
| best_score | 0.821763 | 0.743396 | 0.644092 | 8.136020e-01 | 0.740496 | 0.725935 | 8.683287e-01 |
| lr | 0.000103 | 0.000769 | 0.000329 | 7.281696e-04 | 0.001752 | 0.000557 | 2.479390e-04 |
| weight_decay | 0.000001 | 0.000001 | 0.000004 | 1.376365e-07 | 0.000006 | 0.000001 | 2.406031e-07 |

Table S11: Best hyperparameters of CoxBoost across cancers

| Hyperparameters | BRCA | CESC | HNSC | SARC | UCEC | COAD _ESCA _READ _STAD | GBM _LGG |
|---|---|---|---|---|---|---|---|
| best_score | 0.754921 | 0.741509 | 0.614553 | 0.738693 | 0.735974 | 0.599125 | 0.838529 |
| learning_rate | 0.042043 | 0.04906 | 0.02924 | 0.213336 | 0.273679 | 0.27512 | 0.108908 |
| max_depth | 7 | 6 | 6 | 3 | 6 | 3 | 3 |
| max_features | sqrt | sqrt | sqrt | sqrt | sqrt | sqrt | sqrt |
| min_samples_leaf | 7 | 3 | 8 | 21 | 19 | 9 | 21 |
| min_samples_split | 5 | 8 | 18 | 8 | 16 | 6 | 14 |
| n_estimators | 80 | 98 | 240 | 164 | 74 | 223 | 78 |
| subsample | 0.703887 | 0.821635 | 0.645642 | 0.769202 | 0.996368 | 0.802458 | 0.754964 |

Table S12: Best hyperparameters of Coxnet across cancers

| Hyperparameters | BRCA | CESC | HNSC | SARC | UCEC | COAD _ESCA _READ _STAD | GBM _LGG |
|---|---|---|---|---|---|---|---|
| best_score | 0.812008 | 0.769811 | 0.611671 | 0.811558 | 0.691419 | 0.524202 | 0.645534 |
| l1_ratio | 0.995983 | 0.311345 | 0.214332 | 0.001399 | 0.007990 | 0.433012 | 0.987511 |

Table S13: Best hyperparameters of Cox-nnet across cancers

| Hyperparameters | BRCA | CESC | HNSC | SARC | UCEC | COAD _ESCA _READ _STAD | GBM _LGG |
|---|---|---|---|---|---|---|---|
| best_score | 7.401575e-01 | 0.686792 | 0.481988 | 0.788945 | 0.676568 | 0.660402 | 0.758669 |
| batch_size | 1.600000e+01 | 16.000000 | 24.000000 | 16.000000 | 24.000000 | 24.000000 | 8.000000 |
| dropout_rate | 2.862788e-01 | 0.215288 | 0.174491 | 0.294369 | 0.122659 | 0.145464 | 0.089400 |
| epochs | 2.000000e+02 | 200.000000 | 200.000000 | 200.000000 | 200.000000 | 200.000000 | 200.000000 |
| lr | 5.686883e-04 | 0.004989 | 0.004947 | 0.004902 | 0.000580 | 0.001815 | 0.004645 |
| weight_decay | 1.140310e-07 | 0.000024 | 0.000012 | 0.000016 | 0.000025 | 0.000004 | 0.000002 |

Table S14: Best hyperparameters of DCAP across cancers

| Hyperparameters | BRCA | CESC | HNSC | SARC | UCEC | COAD _ESCA _READ _STAD | GBM _LGG |
|---|---|---|---|---|---|---|---|
| best_score | 0.828248 | 0.803774 | 0.639769 | 0.859296 | 0.716172 | 0.722194 | 0.820315 |
| batch_size | 16.000000 | 16.000000 | 24.000000 | 16.000000 | 24.000000 | 24.000000 | 8.000000 |
| dropout | 0.167603 | 0.019843 | 0.085913 | 0.038519 | 0.154270 | 0.009830 | 0.000113 |
| epochs | 100.000000 | 100.000000 | 100.000000 | 100.000000 | 100.000000 | 100.000000 | 100.000000 |
| lr | 0.004886 | 0.000327 | 0.001752 | 0.000259 | 0.001015 | 0.000851 | 0.001752 |
| noise | 0.267056 | 0.080564 | 0.114802 | 0.141241 | 0.013935 | 0.170603 | 0.166935 |
| weight_decay | 0.000000 | 0.000000 | 0.000000 | 0.000000 | 0.000000 | 0.000000 | 0.000000 |

Table S15: Best hyperparameters of DeepHit across cancers

| Hyperparameters | BRCA | CESC | HNSC | SARC | UCEC | COAD _ESCA _READ _STAD | GBM _LGG |
|---|---|---|---|---|---|---|---|
| best_score | 0.686024 | 0.713208 | 0.682997 | 0.748744 | 0.714521 | 0.69207 | 0.687566 |
| alpha | 0.931403 | 0.598599 | 0.850615 | 0.637459 | 0.90487 | 0.649244 | 0.791605 |
| batch_size | 16 | 16 | 24 | 16 | 24 | 24 | 8 |
| dropout | 0.156714 | 0.137529 | 0.164785 | 0.293165 | 0.25619 | 0.153038 | 0.264258 |
| epochs | 200 | 200 | 200 | 200 | 200 | 200 | 200 |
| hidden_layers | [64, 32, 128] | [256, 32, 256] | [32, 32] | [64, 256] | [64, 128] | [128, 256] | [64] |
| hidden_size_0 | 64 | 256 | 32 | 64 | 64 | 128 | 64 |
| hidden_size_1 | 32.0 | 32.0 | 32.0 | 256.0 | 128.0 | 256.0 | NaN |
| hidden_size_2 | 128.0 | 256.0 | NaN | NaN | NaN | NaN | NaN |
| lr | 0.000184 | 0.000211 | 0.000349 | 0.001272 | 0.002555 | 0.000129 | 0.001524 |
| num_layers | 3 | 3 | 2 | 2 | 2 | 2 | 1 |
| out_features | 20 | 10 | 30 | 10 | 120 | 20 | 120 |
| sigma | 0.01186 | 0.023781 | 0.086834 | 0.024053 | 0.001124 | 0.063037 | 0.008641 |
| weight_decay | 0.000004 | 0.0 | 0.000003 | 0.000001 | 0.000002 | 0.0 | 0.0 |

Table S16: Best hyperparameters of DeepSurv across cancers

| Hyperparameters | BRCA | CESC | HNSC | SARC | UCEC | COAD _ESCA _READ _STAD | GBM _LGG |
|---|---|---|---|---|---|---|---|
| best_score | 0.676181 | 0.788679 | 0.647695 | 0.48995 | 0.707921 | 0.571833 | 0.641681 |
| batch_size | 16 | 16 | 24 | 16 | 24 | 24 | 8 |
| dropout | 0.015838 | 0.219024 | 0.233182 | 0.188284 | 0.190058 | 0.15858 | 0.016385 |
| epochs | 200 | 200 | 200 | 200 | 200 | 200 | 200 |
| hidden_layers | [32, 64] | [128, 128] | [32, 128, 64] | [32, 256, 64] | [128, 32] | [128, 64] | [128, 128] |
| hidden_size_0 | 32 | 128 | 32 | 32 | 128 | 128 | 128 |
| hidden_size_1 | 64 | 128 | 128 | 256 | 32 | 64 | 128 |
| hidden_size_2 | NaN | NaN | 64.0 | 64.0 | NaN | NaN | NaN |
| lr | 0.000428 | 0.000398 | 0.000107 | 0.0001 | 0.004951 | 0.001731 | 0.000164 |
| num_layers | 2 | 2 | 3 | 3 | 2 | 2 | 2 |
| weight_decay | 0.000002 | 0.000002 | 0.0 | 0.000003 | 0.000003 | 0.000009 | 0.00001 |

Table S17: Best hyperparameters of RSF across cancers

| Hyperparameters | BRCA | CESC | HNSC | SARC | UCEC | COAD _ESCA _READ _STAD | GBM _LGG |
|---|---|---|---|---|---|---|---|
| best_score | 0.758366 | 0.724528 | 0.583573 | 0.781407 | 0.734323 | 0.678424 | 0.828021 |
| max_depth | 6 | 7 | 6 | 4 | 7 | 7 | 5 |
| min_samples_leaf | 25 | 22 | 3 | 11 | 6 | 17 | 16 |
| min_samples_split | 14 | 4 | 9 | 3 | 5 | 16 | 9 |
| n_estimators | 121 | 209 | 119 | 363 | 212 | 569 | 212 |

Table S18: Best hyperparameters of Trans-STG across cancers

| Hyperparameters | BRCA | CESC | HNSC | SARC | UCEC | COAD _ESCA _READ _STAD | GBM _LGG |
|---|---|---|---|---|---|---|---|
| best_score | 0.800197 | 0.837736 | 0.706052 | 0.79397 | 0.844884 | 0.717559 | 0.87986 |
| batch_size | 16 | 16 | 24 | 16 | 24 | 24 | 8 |
| d_model | 8 | 16 | 16 | 16 | 8 | 16 | 16 |
| dim_feedforward | 32 | 32 | 8 | 32 | 8 | 8 | 8 |
| dropout | 0.0851292 | 0.0340481 | 0.00950718 | 0.0467148 | 0.0687669 | 0.0994098 | 0.00693402 |
| epochs | 200 | 200 | 200 | 200 | 200 | 200 | 200 |
| lr | 0.000109483 | 0.000653411 | 0.000568071 | 0.000421422 | 0.00346371 | 0.000458326 | 0.000152026 |
| n_heads | 2 | 1 | 4 | 2 | 8 | 2 | 2 |
| n_layers | 1 | 4 | 1 | 2 | 2 | 3 | 3 |
| num_time_bins | 20 | 90 | 120 | 50 | 30 | 30 | 10 |
| weight_decay | 1.06965e-07 | 8.79884e-06 | 7.10758e-05 | 2.09049e-06 | 3.3555e-07 | 3.76847e-05 | 2.57005e-05 |

Table S19: Best hyperparameters of SurvTRACE across cancers

| Hyperparameters | BRCA | CESC | HNSC | SARC | UCEC | COAD _ESCA _READ _STAD | GBM _LGG |
|---|---|---|---|---|---|---|---|
| best_score | 0.807087 | 0.867925 | 0.649856 | 0.836683 | 0.844884 | 0.715757 | 0.860595 |
| attention_dropout | 0.0224406 | 0.0766861 | 0.0786058 | 0.0358624 | 0.0476048 | 0.0377344 | 0.0125565 |
| batch_size | 16 | 16 | 24 | 16 | 24 | 24 | 8 |
| epochs | 200 | 200 | 200 | 200 | 200 | 200 | 200 |
| hidden_dropout | 0.0763616 | 0.0636338 | 0.10668 | 0.266337 | 0.0122195 | 0.161782 | 0.0144127 |
| hidden_size | 8 | 16 | 16 | 16 | 16 | 8 | 16 |
| intermediate_size | 16 | 8 | 64 | 8 | 32 | 16 | 8 |
| lr | 0.00176068 | 0.000270076 | 0.000174012 | 0.000669367 | 0.000380824 | 0.00227691 | 0.000894517 |
| num_heads | 2 | 2 | 2 | 1 | 2 | 2 | 1 |
| num_durations | 50 | 120 | 20 | 20 | 30 | 90 | 50 |
| num_hidden_layers | 1 | 2 | 1 | 1 | 1 | 2 | 2 |
| weight_decay | 6.44734e-05 | 8.13339e-07 | 6.36711e-07 | 1.26919e-05 | 9.4674e-07 | 4.03976e-07 | 2.2904e-07 |

Table S20: Best hyperparameters of Transformer Survival across cancers

| Hyperparameters | BRCA | CESC | HNSC | SARC | UCEC | COAD _ESCA _READ _STAD | GBM _LGG |
|---|---|---|---|---|---|---|---|
| best_score | 0.754183 | 0.856604 | 0.678674 | 0.806533 | 0.839934 | 0.690525 | 0.744308 |
| T_max | 50 | 30 | 10 | 10 | 30 | 50 | 120 |
| alpha | 1 | 1 | 0.1 | 0.1 | 0.5 | 0.5 | 1 |
| batch_size | 16 | 16 | 24 | 16 | 24 | 24 | 8 |
| d_model | 32 | 128 | 128 | 64 | 64 | 128 | 64 |
| dropout | 0.21866 | 0.24803 | 0.00958298 | 0.0741388 | 0.255626 | 0.0902635 | 0.0394987 |
| epochs | 200 | 200 | 200 | 200 | 200 | 200 | 200 |
| ff_model | 256 | 256 | 256 | 128 | 128 | 256 | 512 |
| lr | 0.000563374 | 0.00157532 | 0.000395509 | 0.00333269 | 0.000652544 | 0.000267775 | 0.000227006 |
| nhead | 8 | 4 | 2 | 2 | 8 | 1 | 8 |
| num_layers | 2 | 4 | 3 | 2 | 1 | 4 | 3 |
| weight_decay | 4.40966e-06 | 1.13856e-07 | 1.96738e-07 | 6.39507e-06 | 8.02291e-07 | 3.10274e-06 | 7.9478e-07 |
